# Rat mediodorsal thalamic subdivisions differentially modulate the sensory and affective components of pain through distinct prefrontal pathways

**DOI:** 10.64898/2026.07.17.739109

**Authors:** Hanane Iben-Daoudi, Saadia Ba-M’hamed, Fatima-Zahra Lamghari Moubarrad, Mohamed Bennis, Marc Landry, Zakaria Ouhaz

**Author notes:** Corresponding author: Marc Landry. University of Bordeaux, Centre Broca-Nouvelle Aquitaine, Institute of Neurodegenerative Diseases, IMN, UMR 5293, 146, rue Léo Saignat, Bordeaux, France. share seniority (Marc Landry, Zakaria Ouhaz).

## Abstract

The mediodorsal thalamus (MD) modulates pain through thalamocortical regulation of the mPFC. Yet, MD is often treated as a single anatomical and functional entity despite marked internal heterogeneity. Here, we tested whether medial-central MD (MDmc) and lateral MD (MDl) subdivisions exert dissociable control over sensory-discriminative and affective-motivational components of pain by engaging the anterior cingulate cortex (ACC) and prelimbic cortex (PrL). Using subdivision-selective excitotoxic lesions in rats, combined with anterograde tracing, laminar activity mapping, and projection-specific optogenetic manipulation of MDmc and MDl terminals in ACC or PrL, we determined the contribution of each subdivision and the underlying MD-PFC circuit mechanisms. Behaviorally, MDmc and MDl lesions induced mechanical and thermal hypersensitivity, but only MDmc lesions increased pain-related avoidance. Anatomical analyses showed that MDl preferentially innervated ACC PV cells, whereas MDmc more strongly targeted ACC SOM cells. Lesions further produced subdivision-dependent reorganization of nociception-evoked cFos activity in layers 2/3 and 5 and altered PV/SOM interneuron recruitment. Optogenetic manipulations revealed pathway-specific effects: MDmc-ACC/PrL manipulations enhanced nociceptive gain and avoidance, whereas MDl-ACC inhibition increased hypersensitivity while reducing avoidance, and MDl-PrL inhibition increased both nociceptive sensitivity and avoidance. Together, these findings identify MD subdivision-specific thalamocortical pathways that recruit distinct inhibitory microcircuits within ACC and PrL, thereby differentially shaping sensory-discriminative and affective-motivational components of pain.

## Introduction

Pain is a multidimensional experience in which sensory-discriminative signals are integrated with affective-motivational and cognitive processes that determine threat appraisal, coping, and action selection [50,61,70]. Consequently, nociceptive sensitivity and pain aversiveness can be partially dissociable, such that similar sensory drive may yield different motivational and behavioral outcomes depending on how higher-order circuits assign value and regulate responses [61,70,5]. Within pain-control networks, the medial prefrontal cortex (mPFC) serves as a key integrative hub linking nociceptive information to state evaluation, affect and behavior [51,65]. Its subregions are functionally specialized, with the anterior cingulate cortex (ACC) closely associated with pain aversion, salience processing, and adaptive responses under threat [6,64,28], whereas the prelimbic cortex (PrL) contributes to contextual regulation, executive control, and top-down modulation of nociceptive gain and pain-related behavior [75,74,51,76,44,82]. However, how afferent pathways selectively bias ACC versus PrL processing, and through which local microcircuit interfaces they act, remains incompletely defined [65,2].

Higher-order thalamocortical projections are strong candidates because they regulate cortical excitability and information flow through cortico-thalamo-cortical loops [66,21,77]. We therefore focused on the mediodorsal thalamus (MD), a major higher-order thalamic input to the mPFC. The MD sends dense projections to medial prefrontal territories, including ACC and PrL [33,3,38,36], and is positioned to integrate nociceptive signals through ascending spinal and brainstem inputs [18,40,80,74,26]. Manipulations of medial thalamic regions modulate nociceptive responses in ACC neurons [11,23,22], and recent causal studies indicate that MD-prefrontal coupling can shape both pain-related aversion and sensory hypersensitivity [49,48]. However, the MD is still often treated as a unitary structure, despite evidence that its medial/central (MDmc) and lateral (MDl) territories differ in cytoarchitecture, cortical affiliation, and functional organization [19,39,73,52,21]. Whether these subdivisions engage distinct ACC and PrL circuits to control sensory-discriminative and affective-motivational dimensions of pain differentially remains unknown.

Long-range thalamic inputs interact with layer-specific cortical architecture to shape integration and output in distinct ways, with superficial versus deep layers supporting different computational roles and output channels [2]. In parallel, inhibitory interneuron classes provide complementary control motifs. Parvalbumin (PV) interneurons can impose fast, perisomatic gain control and feedforward inhibition, whereas somatostatin (SOM) interneurons preferentially regulate dendritic integration and context-dependent gating of pyramidal activity [71]. Notably, MD-ACC signaling has been shown to recruit feedforward inhibition through local interneuron circuits, providing a concrete entry point by which MD channels can regulate prefrontal gain and timing [8,49,46]. Together, these principles motivate the central question of this study: whether MD subdivision identity (MDmc vs MDl) and prefrontal target (ACC vs PrL) specify distinct laminar and PV/SOM microcircuit engagement that can explain differential control over pain dimensions.

Here, we directly test the hypothesis that MDmc and MDl exert dissociable control over distinct pain components by engaging ACC versus PrL through layer- and interneuron-specific mechanisms. We combine subdivision-selective lesions, pathway-resolved anatomical tracing, laminar and interneuron-specific activity readouts, and projection-specific optogenetic inhibition and activation to link MD subdivision organization to ACC/PrL microcircuit engagement and causal control of sensory-discriminative and affective-motivational components of pain.

## Materials and Methods

### 1. Animals

A total of 131 adult male and female Sprague-Dawley rats (250-400 g at surgery) were obtained from the EOPS facility (Broca center animal facility, University of Bordeaux) and Janvier Labs (Le Genest-Saint-Isle, France). Animals were housed 2–3 per cage under controlled conditions (22-23 °C, 40% relative humidity, 12 h light/dark illumination cycle; lights on at 07:00) with food and water available ad libitum. Rats were acclimatized to laboratory conditions for at least one week before experiments. All procedures were approved by the local ethics committee and the French Ministry of Higher Education, Research and Innovation (APAFIS#46446) and complied with applicable national and European directive 2010-63-EU. Humane care was provided throughout, and every effort was made to minimize distress. Animals were randomly assigned to experimental groups, and behavioral testing, image acquisition, histological quantification, and statistical analyses were performed by experimenters blinded to group allocation whenever possible.

### 2. Surgery

#### 2.1. Pharmacological lesioning

In the lesion studies, rats received buprenorphine (0.05 mg/kg) subcutaneously fifteen minutes before anesthesia. Anesthetized rats (5 mg/kg xylazine, 50 mg/kg ketamine mixture, i.p.) were fixed in a David-Kopf stereotaxic frame in a flat skull position. Microinfusions of 0.12 M N-methyl-d-aspartate (NMDA, Sigma Aldrich, USA) dissolved in phosphate buffer (pH 7.20) were administered bilaterally via a 1 µl Hamilton syringe over 5 minutes in the regions of interest: the MDmc and the MDl subdivisions (8 males and 8 females per group), following a published protocol [24]. Subdivisions of the MD were targeted using Paxinos and Watson coordinates (mm from bregma). MDmc received two injections per hemisphere, one anterior (AP -2.8, ML +0.2, DV -5.4, delivering 0.15 µL) and one posterior (AP -3.2, ML +0.2, DV -5.5, delivering 0.15 µL). In contrast, MDl received a single injection (AP -3.5, ML +0.7, DV -5.5, delivering 0.10 µL). After each injection, the needle remained in place for 10 minutes to facilitate diffusion and minimize NMDA wicking. Using the same injection coordinates, the sham group received injections of sterile phosphate buffer instead of NMDA. After surgery, rats were individually housed for up to five days during recovery and then returned to their pre-surgery housing cohort.

#### 2.2. Viral constructs and injections

For optogenetic manipulation, adeno-associated viral (AAV) vectors were used under the CaMKIIα promoter. For optogenetic activation, we injected AAV5-CaMKIIα-hChR2(H134R)-EYFP (Addgene #26969-AAV5; titer ≥1×10¹³ vg/mL). For inhibition we used AAV5-CaMKIIα-ArchT-GFP (Addgene #99039-AAV5; titer ≥7×10¹² vg/mL). Control animals received AAV5-CaMKIIα-EGFP (Addgene #50469-AAV5; titer ≥3×10¹² vg/mL). Rats were anesthetized with isoflurane (5% for induction and 1.5-2% for maintenance) and placed in a stereotaxic frame (5-6 males and females per optogenetic group). Pre-emptive analgesia consisted of buprenorphine 0.05 mg/kg SC and local infiltration of lidocaine 2 mg/kg at the incision site. Using the same coordinates in the lesion study, MDmc groups received two injections, one anterior (delivering 0.25 µL) and one posterior (delivering 0.20 µL) to cover the rostro-caudal extent of the medial magnocellular sector. MDl groups received a single injection (delivering 0.15 µL). Virus was infused at 0.05 µL/min through a pulled-glass micropipette coupled to a nanoliter injector. The pipette remained in place for 10 min before withdrawal to limit reflux.

For anterograde anatomical tracing of the MD subdivision-PFC pathway, we injected AAV5-CaMKIIα-EGFP unilaterally using the same stereotaxic approach and the same MD coordinates/volumes as above (two injections into the MDmc and a single injection into the MDl). Animals survived 4 weeks to allow expression and axonal transport, then were perfused. Only male groups were used, with 5 rats in the MDmc group and 6 rats in the MDl group. Anatomical tracing was performed in males only to limit animal use and because this experiment aimed to define projection architecture rather than sex-dependent behavioral modulation. Sex was nevertheless included as a biological variable in lesion, optogenetic, and activity-mapping experiments.

#### 2.3. Optic fiber implantation and projection targeting

To stimulate MD terminals in the PFC, optical cannula (LC Ceramic Ferule NO flange OD 1.25 mm and ID 200 µm, associated with Ø200 µm Core TECS-Clad Multimode Optical Fiber, 0.5 NA) was implanted ipsilateral to the MD injection during the same surgery, either above the ACC (AP +2.9, ML +0.6, DV -2.5) or the PrL cortex (AP +3.3, ML +0.6, DV -3.1). Optic fiber efficiency was tested beforehand and was used if it reached a minimum efficiency of 90%. The surface of the prefrontal area was cleaned with hydrogen peroxide and water, a burr hole was drilled with a handheld drill, and the optic fiber was slowly inserted into the target (PrL or ACC at the coordinates above). A thin layer of C&B Metabond adhesive was applied to the skull, then dental cement was added to secure the fiberoptic cannula. Incisions were closed with sutures, and animals recovered on a heated pad. Meloxicam 5 mg/kg s.c. was administered once daily for 4 post-operative days, and animals were housed singly during this period. Behavioral experiments began four weeks after surgery to ensure robust expression and anterograde transport.

### 3. Optogenetic stimulation protocols

Before testing, animals were handled daily for 5-6 days (about 6-10 min/day) to habituate to human handling. On each test day, animals were transported to the testing facility at least 2 h before experimentation. In the presence of a brief 0.1 mL isoflurane exposure, the implanted ferrule was connected to a patch cable (200 µm core, 0.5 NA, multimode) using a 2.5-mm zirconia mating sleeve; the patch cable was then connected to the laser generator via a rotary joint. Animals were placed in small enclosed testing arenas, with cable tension minimized to allow free mobility for ∼60 min before trials. Optical power was measured before each experiment using a power sensor. Optical stimulation was delivered through a 473 nm laser for activation (ChR2) or a 593 nm laser for inhibition (ArchT). Light trains during behavior consisted of 10-ms pulses at 20 Hz for activation or continuous light for inhibition, with output power calibrated at the fiber tip (5-8 mW for terminals).

### 4. Behavioral assays

#### 4.1. Mechanical sensitivity

The paw withdrawal threshold was measured by applying a gradually increasing pressure stimulus to the plantar surface of the hind paw, assessing the nociceptive response to mechanical stimulation [55,9]. Rats were individually placed in a transparent plastic box with penetrable metal mesh bottoms (5 x 5 mm squares) and habituated to the test environment for 60 min before testing procedures. A series of von Frey filaments ranging in size from 4 to 300 g was applied perpendicularly to the plantar surface of the right and left hind legs, in a consecutive ascending manner. Each filament is applied seven times for three to five seconds until it buckles. The positive reaction is characterized by noxious behavior such as quick withdrawal or licking of the paw, which occurs either during stimulus administration or immediately after removal of the filament. In general, the filament reaches the nociception threshold if it generates at least four of the seven painful reactions. For optogenetic testing, animals were tested individually before, during, and after the lights were turned on, with a 1-hour interval between measurements. Both ipsilateral (ipsiL) and contralateral (ContraL) paws were tested. The light was turned on 3 minutes before the measurements began and kept on until all the measurements were finished.

#### 4.2. Thermal sensitivity

The hot/cold plate was used to test the sensitivity to thermal pain by evaluating paw withdrawal latency to nociceptive thermal stimuli [13]. The test takes place on a rectangular (20cm) metal heating plate, topped by a transparent polyester cylinder (20cm diameter, 25cm high). During the test, the plate is maintained at a constant temperature (5°C for the cold plate and 55°C for the hot plate), and the animal is placed in free movement in synchronization with the timer’s triggering and light activation for optogenetic conditions. The exposure time is a maximum duration of 30 seconds, which favors the expression of various noxious behaviors (withdrawal, licking of the paws, or jumping) and respects the ethical guidelines by preventing tissue damage. The day of thermal evaluation was different from the day of mechanical evaluation for each animal.

#### 4.3. Place escape/avoidance paradigm

The place escape/avoidance paradigm (PEAP) is used to study pain’s affective and emotional components independently of the other components. The test is based on the inhibition of innate or acquired behaviors by association with aversive stimulation [41,42]. The behavior investigated here is the natural tendency of rodents to abandon wide, brightly lit spaces and hide in confined, dark ones, using mechanical stimulation by von Frey filaments as an aversive stimulus. The apparatus consists of two compartments of equal dimensions (30 x 30 x 60 cm, each), without top and bottom, one lighted and the other dark, positioned on a mesh screen. The protocol consists of two phases: a first habituation phase, in which the animal is confined for 10 min in each compartment, and a second test session phase, in which the rat is free to move around for 30 minutes. When the animal is located within the dark compartment, the right paw is stimulated by a suprathreshold von Frey filament, and when it moves to the lighted compartment receives an infra-threshold stimulation on the same paw at 15 s intervals. For the optogenetic cohort, some animals were tested with light off (MDmc/light off and MDl/light off group), other animals were tested with light on (MDmc/light on and MDl/light on group) (yellow or blue laser). The location of the animal (within the light side or dark side) was recorded and converted to a percentage of time spent in the light side of the chamber.

### 5. Immunohistochemistry

Animals were deeply anesthetized with xylazine (5 mg/kg, i.p.) and ketamine (50 mg/kg, i.p.), then transcardially perfused with saline followed by 4% paraformaldehyde (PFA) in 0.1 M PBS. Brains were post-fixed overnight in 4% PFA, transferred to PBS, and sectioned coronally on a vibratome (Leica VT 1000 S). Free-floating sections were rinsed three times for 10 min in 0.1 M PBS, incubated for 1 h at room temperature in a blocking solution of PBS containing 1% BSA and 0.3% Triton X-100, then placed in primary antibody solution prepared in the same buffer and incubated overnight at 4 °C. After three additional 10-min PBS washes, sections were incubated for 2 h at room temperature with fluorophore-conjugated secondary antibodies diluted 1:500 in blocking buffer, washed again, mounted on Superfrost slides, and coverslipped with Fluoromount-G. Negative controls were performed by omitting the primary antibodies, which abolished specific staining. Antibody specificity was further supported by the manufacturers’ validation data and previous use in rat brain tissue.

Verification of viral expression and injection specificity in the MD used Nissl staining or GFP immunofluorescence on 40 µm sections through the MD. GFP labeling was performed with chicken anti-GFP (1:500, Aves Labs, GFP-1010) followed by goat anti-chicken Alexa Fluor 488 (1:500, Invitrogen, A78948). Only animals with confined expression to MDmc or MDl were retained for analysis. Cases with off-target spread into midline or intralaminar nuclei were excluded a priori.

To confirm fiber-optic placement and light-evoked neuronal activation in PFC during behavioral testing, we performed cFos immunostaining on 40 µm sections from the anterior block containing ACC and PrL cortex. Animals (n = 5/6 per group) were exposed to the same environment and the same laser parameters used in the von Frey sessions, brains were collected 3 h later to allow cFos protein accumulation, and sections were processed with rabbit anti-cFos (1:500, Synaptic systems, 226008) and chicken anti-GFP followed by Alexa Fluor 568 goat anti-rabbit secondary for cFos (1:500, Invitrogen, A11011) and Alexa Fluor 488 donkey anti chicken for GFP.

For laminar mapping of activity after MD lesions, we quantified cFos, cFos /PV, and cFos /SOM in the ACC and PrL cortex (n = 6/7 males and females per group). Free-floating sections (40 µm) were processed with rabbit anti-cFos (1:500, Synaptic Systems #226 008) and chicken anti-PV (1:3000, Novus Biologicals NBP2-50036SS), or rabbit anti-cFos (1:500) and guinea pig anti-SOM (1:1000, Synaptic Systems #366 004). Primaries were followed with fluorophore-conjugated secondaries according to the same protocol shared above (all 1:500): Alexa Fluor 568 goat anti-rabbit for cFos, Alexa Fluor 488 donkey anti-chicken for PV, and Alexa Fluor 488 goat anti-guinea pig (Invitrogen A11073) for SOM. After final washes, sections were mounted with Fluoromount-G.

For anterograde tracing and interneuron target identification, 20 µm coronal sections through ACC and PrL cortex were triple-labeled for GFP, PV, and SOM using the same protocol. Primary antibodies were guinea pig anti-PV (1:3000, Synaptic Systems, 195 308), rabbit anti-SOM (1:1000, Novus Biologicals NBP1-87022), and chicken anti-GFP (1:500). Corresponding secondaries were Alexa Fluor 488 donkey anti-chicken (1:500) for GFP, Alexa Fluor 555+ donkey anti-rabbit (1:500, Invitrogen A32794) for SOM, and Alexa Fluor 647 goat anti-guinea pig (1:500, Invitrogen A21450) for PV.

### 6. Histological evaluation of the lesion site

Brains were removed and post-fixed in PFA overnight at 4°C. Following post-fixation, coronal sections were cut at 40µm using a vibratome (Leica VT1200) and mounted onto gelatin-coated slides. Slides were air-dried and subsequently immersed in a 0.1% cresyl violet solution for 2 minutes. Dehydration was performed through a graded series of ethanol solutions (50%, 70%, and 100% ethanol for 2 minutes each), followed by clearing in xylene (2 minutes) and cover-slipped with Eukitt mounting medium. Lesion boundaries were then visualized using a light microscope (Olympus Corporation Model CX21FS1) and mapped onto corresponding sections of the rat brain atlas of Paxinos and Watson (2008) to confirm accurate targeting. In cases where lesions extended beyond the intended target area, animals were excluded from further analysis. Neurons were identified based on their larger soma size, prominent Nissl substance, and clear nuclear morphology, including a visible nucleolus, which distinguishes them from smaller glial cells that lack such cytoplasmic granularity and exhibit smaller, more densely stained nuclei [17].

### 7. Image acquisition and analysis

For verification of viral expression in the injection site, images were taken at 20x with an epifluorescence microscope (Olympus BX63). GFP expression in MD was assessed; animals with off-target MD expression (intralaminar/midline nuclei) or misplaced fibers were excluded a priori according to predefined exclusion criteria. This resulted in the following exclusions: insufficient unilateral viral expression: two MDmc-ACC- ChR2, one MDl-ACC-ChR2, one MDl-ACC-ArchT, one MDl-PrL-ChR2 rats; misplaced MD target: two MDmc-PrL-ArchT, two MDmc-PrL-ChR2, and one MDl-ACC-ChR2 rats.

For optical fiber placement verification and cFos/interneurons activity mapping in behavioral experiments, brain slices were imaged using a slide scanner (Nanozoomer 2.0HT, Hamamatsu Photonics, Hamamatsu, Japan) at low magnification to capture the full coronal section. Sections were acquired with identical exposure settings across animals. Quantification was performed in ImageJ: ACC/PrL ROIs were delineated using atlas landmarks.

For anterograde tracing, image stacks of the ACC and the PrL cortex were acquired using a confocal microscope (Leica TCS SP5), with a 20x oil immersion objective (z-step = 1 µm; stack depth = 20 µm). For each animal and region, three non-overlapping fields per hemisphere were sampled in layer-matched ROIs defined using Paxinos and Watson coordinates (5-6 per group as indicated in figure legends). Stacks were analyzed with a semi-automatic quantification with an in-house ImageJ-based macro for co-localization of fluorescent markers present on a focal plane on confocal fluorescent images called BioLoc3D [10]. In BioLoc3D, the GFP channel was assigned as the projection channel and PV or SOM as the target interneuron channel; thresholds and size filters were calibrated on a subset of stacks and then fixed for the entire dataset. The pipeline returned projection volume, surface, mean gray value (MGV), and integrated density within ROIs, as well as putative GFP-PV/SOM contact counts after 3D object reconstruction and distance-based assignment.

### 8. Statistical analyses

Statistical analyses were performed using SigmaPlot 12.5 (SigmaStat; Systat Software Inc., San Jose, CA, USA). Data are presented as mean ± SEM. Assumptions for parametric tests were verified before analysis (normality: Shapiro-Wilk; homogeneity of variance: Levene). Unless otherwise stated, ANOVA models were followed by Holm-Šídák *post-hoc* multiple-comparisons procedures when significant main effects and/or interactions were detected. For lesion experiments, behavioral and immunohistochemical outcomes were analyzed using two-way ANOVA with lesion and sex as factors. For optogenetic sensory assays, measures collected across experimental phases (pre-light, light delivery, post-light) were analyzed using two-way repeated-measures ANOVA with phase as the within-subject factor and sex as the between-subject factor. PEAP performance was analyzed using ordinary three-way ANOVA with optogenetic condition, subdivision, and sex as factors. Percent-normalized behavioral measures (normalized to the light-off or pre-opto baseline) were analyzed using two-way ANOVA, with Holm-Šídák correction for planned comparisons. Tracing outcomes were compared between the MDmc and MDl groups using two-tailed unpaired t-tests (Welch’s correction when variances were unequal) or Mann-Whitney tests when normality assumptions were not met. Statistical significance was set at p < 0.05.

## Results

### 1. MDmc and MDl lesions modulate mechanical and thermal nociception

We verified with Nissl staining that NMDA lesions were mostly restricted to the intended MD subdivisions (MDmc or MDl) across the rostro-caudal axis (Fig. S1A-C). Quantitative analyses demonstrated a marked reduction in neuronal density within the targeted subdivision, with minimal loss in non-targeted MD subregions (Fig. S1D). Off-target damage in neighboring thalamic nuclei was limited, supporting lesion specificity (Fig. S1E).

To test whether MDmc and MDl excitotoxic lesions alter baseline mechanical and thermal pain sensitivity in male and female rats, we quantified paw withdrawal responses to mechanical stimulation (von Frey test), noxious heat (hot plate), and noxious cold (cold plate) (Fig. 1A). Von Frey withdrawal analysis revealed main effects of lesion (*F*_(2,42)_ = 67.43, p < 0.001) and sex (*F*_(1,42)_ = 16.25, p < 0.001) on withdrawal thresholds, but no sex × lesion interaction (*F*_(2,42)_ = 0.84, p = 0.438) (Fig. 1B). In both males and females, von Frey thresholds were significantly reduced in MDmc and MDl animals compared with sham (males: t = 8.64 and 6.87, p < 0.001; females: t = 6.87 and 5.58, p < 0.001).

**Figure 1.**
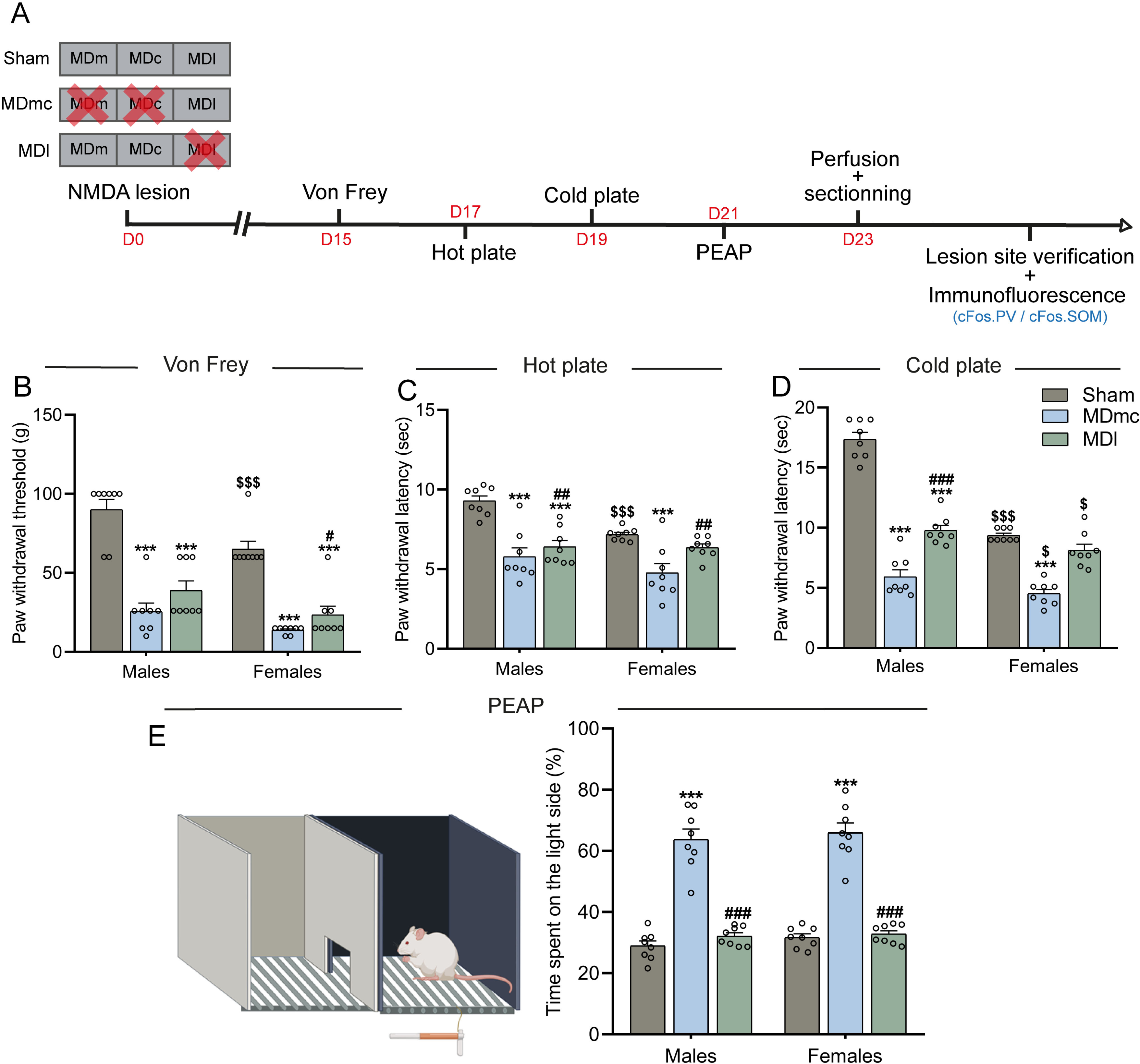
Functional dissociation of MD subdivisions in the sensory-discriminative and affective dimensions of pain. (A) Experimental design of the excitotoxic lesion strategy. (B) Paw withdrawal thresholds (g) in the Von Frey test. (C) Withdrawal latencies (s) in the hot plate. (D) Cold-evoked withdrawal latencies (s) in the cold plate. (E) schematic representation of the PEAP (left). Percentage of time spent in the illuminated compartment of the PEAP (right). Data are presented as mean ± SEM (n = 8 per group). Behavioral data were analyzed using two-way ANOVA with lesion (Sham, MDmc, MDl) and sex (male, female) as factors, followed by Holm-Šídák *post-hoc* tests for multiple comparisons. ***p < 0.001 lesioned animal vs Sham. #p < 0.05, ##p < 0.01, ###p < 0.001 MDmc vs MDl. $p < 0.05, $$p < 0.01, $$$p < 0.001 female vs male.

We then assessed whether MDmc and MDl lesions similarly impact acute thermal (hot and cold) pain responses in the hot/cold plate tests. For the hot plate the analysis revealed main effects of lesion (*F*_(2,42)_ = 26.81, p < 0.001) and sex (*F*_(1,42)_= 10.05, p = 0.003) on withdrawal latencies. In contrast, the lesion × sex interaction did not reach significance (*F*_(2,42)_ = 3.22, p = 0.050) (Fig. 1C). *Post-hoc* comparisons performed within each sex showed that, in males, both MDmc and MDl lesions shortened hot plate latencies relative to sham (t = 6.06; t = 5.00; p < 0.001, respectively); however, no differences were observed between the two lesioned groups (t = 1.06, p = 0.295). In females, MDmc lesions also reduced latencies compared with sham (t = 4.19, p < 0.001). In contrast, the MDl lesion groups did not differ from the sham groups (t = 1.42, p = 0.164). In addition, latencies were longer in the MDl than in the MDmc lesion group (t = 2.77, p = 0.017).

Cold plate latencies showed a more pronounced influence of MD lesions and sex. Two-way ANOVA revealed main effects of lesion (*F*_(2,42)_ = 160.11, p < 0.001), sex (*F*_(1,42)_ = 97.81, p < 0.001), and lesion × sex interaction (*F*_(2,42)_ = 33.77, p < 0.001) on withdrawal latencies (Fig. 1D). In males, both MDmc and MDl lesions markedly shortened withdrawal latencies relative to their sham (t = 17.77; t = 11.78; p < 0.001, respectively), and latencies were shorter in MDmc than in MDl lesioned males (t = 5.99, p < 0.001). In females, MDmc lesions likewise reduced latencies compared with sham (t = 7.51, p < 0.001), and MDl females showed longer latencies than MDmc females (t = 5.59, p < 0.001), whereas sham and MDl females did not differ (t = 1.92, p = 0.062).

### 2. MDmc and MDl lesions differentially shape escape-avoidance responses to mechanical pain

We next asked whether MDmc and MDl influence the affective-motivational component of pain by assessing escape-avoidance behavior in the PEAP.

Performance in the PEAP was strongly affected by the lesion condition. Two-way ANOVA revealed a strong main effect of lesion (*F*_(2,42)_ = 155.48, p < 0.001) on the time spent in the light compartment, whereas neither the main effect of sex (*F*_(1,42)_ = 1.11, p = 0.297) nor the lesion × sex interaction (*F*_(2,42)_ = 0.12, p = 0.887) reached significance (Fig. 1E). MDmc-lesioned groups displayed a marked increase in avoidance of the nociceptive compartment. Both male and female MDmc rats spent more time in the non-noxious (light) side than shams (males: t = 11.22; females: t = 11.04, p < 0.001), and more than MDl-lesioned rats of the same sex (males: t = 10.19; females: t = 10.68, p < 0.001). By contrast, MDl-lesioned groups did not differ from sham control animals in either sex (males: t = 1.03, p = 0.309; females: t = 0.36, p = 0.721).

Together with the von Frey data, these findings indicate that MDmc lesions are associated with enhanced escape-avoidance of the nociceptive compartment, whereas MDl-lesioned rats, despite exhibiting comparable mechanical hypersensitivity, fail to increase avoidance behavior. This dissociation suggests that MDl damage selectively disrupts the affective-motivational response to mechanical pain.

### 3. MDmc and MDl differentially target PV and SOM interneurons in ACC and PrL

#### 3.1. MDl projections form denser arborizations than MDmc in the mPFC

Given the distinct effects of MDmc and MDl lesions on affective pain behaviors, we next asked whether these functional differences are mirrored by a different anatomical organization of their projections onto prefrontal inhibitory microcircuits. Using a GFP-expressing AAV strategy, we quantified the density of MDmc- and MDl projections to the mPFC (Fig. 2A-B). This analysis revealed that MDl projections to the ACC were overall denser than those of MDmc. In the ACC, MDl injections represented a larger GFP⁺ volume than MDmc (p = 0.014) (Fig. 2C-D). MGV measurements further indicated higher GFP signal intensity in the ACC (Fig. 2E) for MDl than for MDmc projections (p< 0.001). A similar effect on GFP⁺ volume was found in the PrL (Fig. 2F-G) (p < 0.001), whereas MGV did not differ between groups in the PrL (p = 0.294) (Fig. 2H-I).

**Figure 2.**
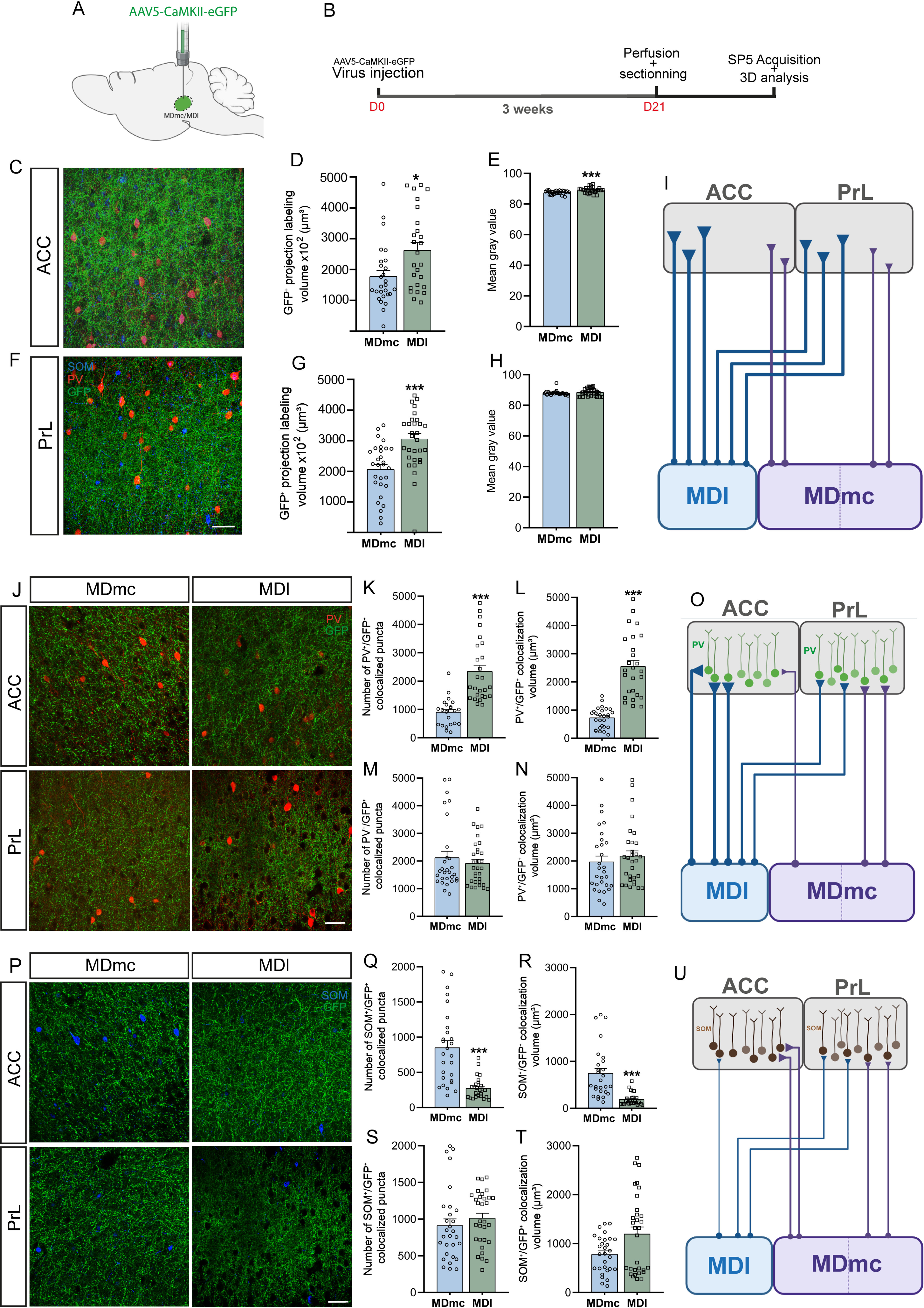
MDmc and MDl projections to ACC and PrL cortex differentially target PV⁺ and SOM⁺ interneurons. (A) Schematic of AAV5-CaMKII-eGFP injection into MDmc/MDl. (B) Experimental timeline. (C-H) Representative 3D confocal stacks and quantification of GFP⁺ projection signal in ACC (C-E) and PrL (F-H) [volume (µm^3^) and mean gray value]. (I) Summary wiring schematic. (J-N) Representative confocal images showing GFP⁺ terminals from MDmc or MDl in close apposition to PV⁺ interneurons in ACC and PrL, with quantification of PV/GFP⁺ colocalized puncta and volume in the ACC and the PrL. (O) Summary schematic of PV targeting. (P-T) Same analysis for SOM⁺ interneurons (representative images and quantification of SOM/GFP⁺ colocalized puncta and volume). (U) Conceptual summary of SOM targeting. Scale bar: 50 µm. Data are presented as mean ± SEM (N = 5-6 animals per group; n = 26-31 mean stacks per section). Comparisons between MDmc and MDl groups were performed using unpaired t-tests with Welch’s correction when variances were unequal, or Mann-Whitney non-parametric tests when distributions were non-normal. *p < 0.05, **p < 0.01, ***p < 0.001, MDmc vs MDl.

#### 3.2. MDl preferentially innervates PV interneurons, whereas MDmc more strongly targets SOM cells

We then asked how these projection patterns map onto specific inhibitory cell types by quantifying colocalization of PV⁺/GFP⁺ and SOM⁺/GFP⁺ in ACC and PrL. In the ACC, MDl projections formed many more PV⁺/GFP⁺ contacts than MDmc projections (p < 0.001) (Fig. 2J-K), and the volume of PV⁺/GFP⁺ colocalization was also markedly larger for MDl than for MDmc (p < 0.001) (Fig. 2L). In contrast, in the PrL, the number of PV⁺/GFP⁺ contacts (p = 0.837) (Fig. 2M) and their volume (p = 0.303) (Fig. 2N) remained similar, indicating similar projections from MDmc and MDl on PV interneurons (Fig. 2O).

The pattern was reversed for SOM cells in ACC (Fig. 2P), where MDmc projections gave rise to more SOM⁺/GFP⁺ contacts than MDl projections (p < 0.001) (Fig. 2Q). Also, the volume of SOM⁺/GFP⁺ colocalization was likewise greater for MDmc than MDl projections (p < 0.001) (Fig. 2R). In the PrL, neither the number (p = 0.132) (Fig. 2S) nor the volume (p = 0.073) of SOM⁺/GFP⁺ colocalization (Fig. 2T) differed between MDmc and MDl. Taken together, these tracing data support a circuit architecture in which MDl projections preferentially engage PV interneurons. In contrast, MDmc projections more strongly recruit SOM interneurons in the ACC, but not in the PrL (Fig. 2U).

### 4. MD lesions trigger specific cFos activity and PV/SOM recruitment in mPFC

We next asked how MD damage reshapes activity-dependent cortical recruitment across layers. We quantified laminar cFos expression as a readout of overall activation following a nociceptive stimulation, and then measured cFos co-expression in PV⁺ and SOM⁺ interneurons (Figs. 3-4; Tables S1-S3). Across both ACC and PrL, cFos activation and its expression in PV/SOM interneurons were not observed in layers 1 and 6.

**Figure 3.**
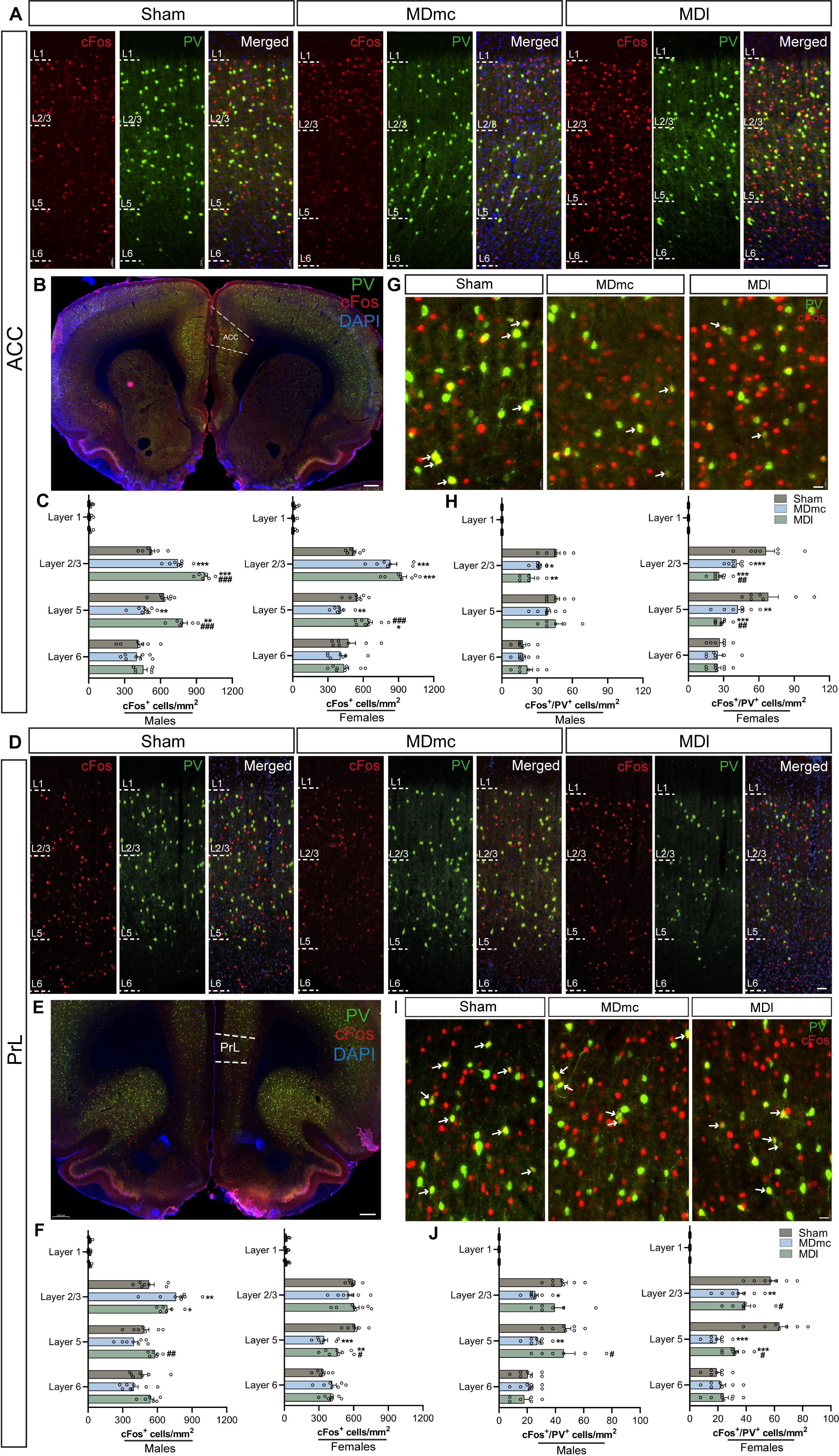
MD subdivision lesions reshape laminar PV interneuron recruitment indexed by cFos in ACC and PrL. ACC **(A-C, G-H)**. **(A)** Representative laminar images of the ACC showing cFos (red), PV (green), and merged channels (including DAPI, blue) across cortical layers in Sham, MDmc, and MDl groups. Sections are oriented with the pial surface at the top and the white matter at the bottom; white dashed lines indicate layer boundaries used for quantification. Scale bar: 50 µm. **(B)** Low-magnification coronal overview showing the ACC ROI used for laminar quantification. Scale bar: 500 µm. **(C)** Quantitative analysis of total cFos⁺ cell density (cells/mm²) in ACC layers in males (left) and females (right). PrL cortex **(D-F, I-J)**. Same workflow for PrL, including laminar (Scale bar: 50 µm) and overview images (Scale bar: 500 µm) **(D-E)**, and layer-resolved densities of total cFos⁺ nuclei **(F)** across cortical layers 1, 2/3, 5, and 6, split by sex. **(G)** High-magnification fields illustrating double-labelled cFos⁺/PV⁺ cells (white arrows). Scale bar: 50 µm. **(H)** Quantitative analysis of double-labelled cFos⁺/PV⁺ interneuron density (cells/mm²) in ACC layers in males (left) and females (right). High-magnification examples (Scale bar: 50 µm) **(I)**, and layer-resolved densities of cFos⁺/PV⁺ interneurons **(J)** across cortical layers 1, 2/3, 5, and 6, split by sex. Data are presented as mean ± SEM. Sham (females n = 7, males n = 7), MDmc (females n = 7, males n = 8), and MDl (females n = 8, males n = 6). For each cortical layer and region, laminar densities were analyzed using two-way ANOVA with factors lesion and sex, followed by Holm-Šidák post-hoc multiple-comparisons tests. *p < 0.05, **p < 0.01, ***p < 0.001 lesioned vs Sham; #p < 0.05, ##p < 0.01 MDl vs MDmc.

**Figure 4.**
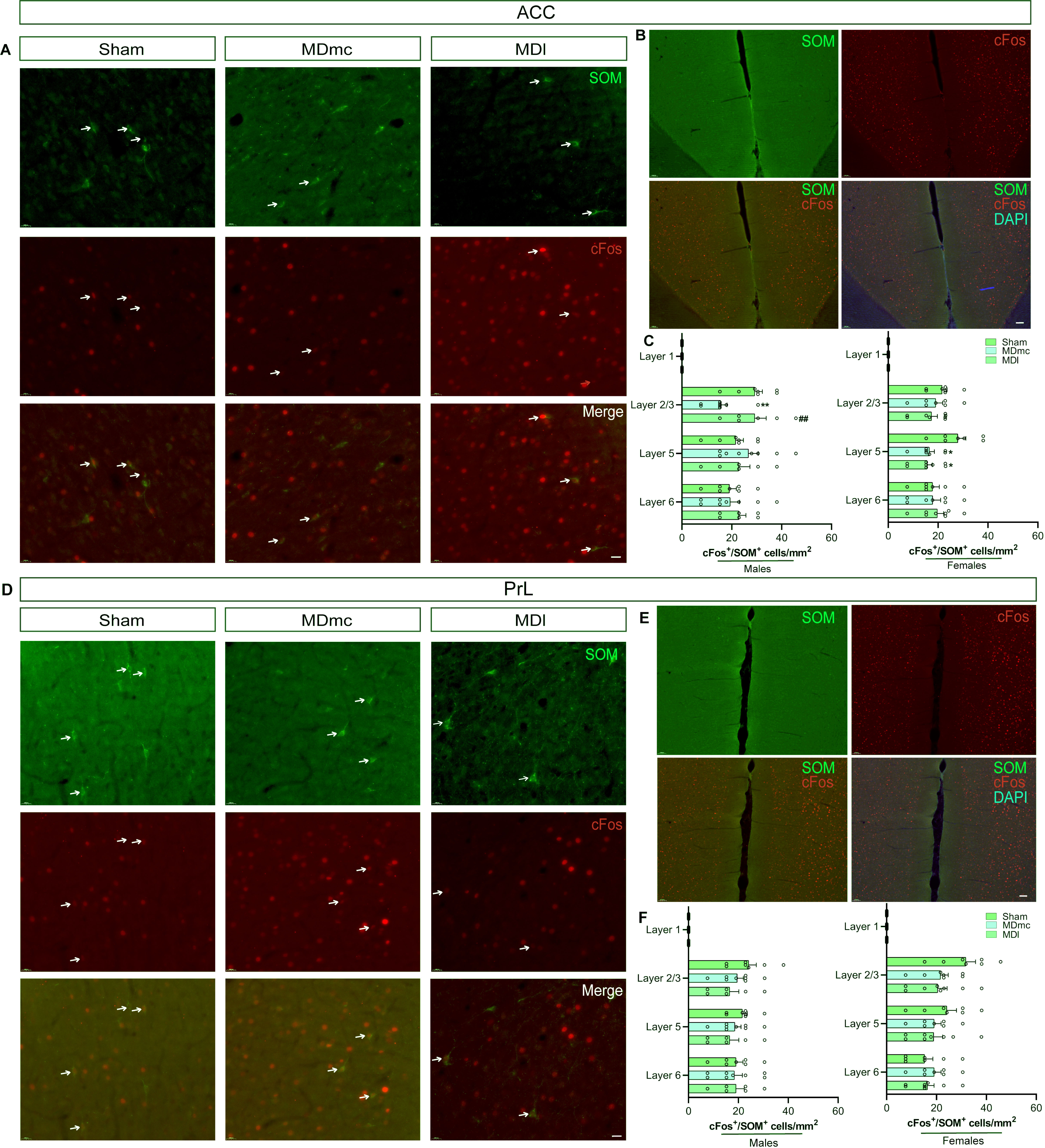
MDmc lesions selectively reduce SOM interneuron activation in ACC. ACC **(A-C). (A)** Representative images of the ACC showing SOM⁺ interneurons (green), cFos⁺ nuclei (red), and merge in sham, MDmc, and MDl groups. Double-labelled cFos⁺/SOM⁺ cells are visible in the merged panels and indicated by arrows. Scale bar: 50 µm. **(B)** Low-magnification coronal overview illustrating the ACC anatomical level used for analysis; SOM^+^ (green), cFos^+^ (red), SOM^+^/cFos^+^ merge, and SOM^+^/cFos^+^/DAPI (blue) merge. Scale bar: 100 µm. **(C)** Quantitative analysis of double-labelled cFos⁺/SOM⁺ interneuron density (cells/mm²) in males (left graph) and females (right graph) across ACC layers. **(D-F)** PrL: Same analysis in PrL, including representative images **(D)**, coronal overview of the PrL ROI **(E)**, and layer-quantification of cFos⁺/SOM⁺ interneuron densities split by sex **(F)**. Data are presented as mean ± SEM. Sham (females n = 7, males n = 7), MDmc (females n = 7, males n = 8), and MDl (females n = 8, males n = 6). For each layer and region, laminar densities were analyzed using two-way ANOVA with factors lesion and sex, followed by Holm-Šidák *post-hoc* multiple-comparisons tests. *p < 0.05, **p < 0.01 lesioned vs Sham; ##p < 0.01 MDl vs MDmc.

#### 4.1. MDmc and MDl lesions impose distinct laminar cFos activation profiles in ACC and PrL

In the ACC, layer 2/3 cFos density increased after both MDmc and MDl lesions relative to sham in males and females (Fig. 3A-B; Table S1). In males, the increase was stronger after MDl lesions than after MDmc lesions. In contrast, females showed similarly elevated cFos levels in the two lesion groups (Fig. 3C). In layer 5, cFos density was differentially modulated, showing a decrease after MDmc lesion, and an increase after MDl lesion in both males and females. MDl lesions produced higher cFos levels than MDmc lesions in both sexes.

In the PrL, lesion effects were again localized to layers 2/3 and 5, with clearer sex dependence (Fig. 3D-E; Table S1). In layer 2/3, cFos density was increased after MDmc and MDl lesions compared with sham in both sexes; additionally, within the MDmc group, males showed higher cFos recruitment than females. In layer 5, MDmc lesions showed a tendency toward reduced cFos recruitment, whereas MDl lesions showed a tendency toward increased recruitment, and this separation was more evident in males (Fig. 3F; Table S1).

Taken together, these results indicate that MDmc and MDl lesions leave layer 1 largely unaffected but robustly enhance cFos recruitment in layers 2/3 and 5 of ACC and PrL, with MDl lesions driving sex-dependent effects in PrL.

#### 4.2. MD lesions reduce PV interneuron activation in a lamina- and subdivision-specific manner

To assess PV interneuron recruitment in response to noxious stimulation, we quantified PV⁺ cells co-expressing cFos (PV⁺/cFos⁺) across layers in ACC and PrL cortices.

In the ACC, layer 2/3 PV recruitment decreased after both MDmc and MDl lesions compared with sham in males and females (Fig. 3G). In females, PV recruitment was lower after MDl lesions than after MDmc lesions (Fig. 3H; Table S2). In layer 5, females again showed clear reductions after both lesions, with MDl females showing the lowest PV recruitment, whereas males showed comparatively modest lesion-related changes (Fig. 3H; Table S2). Thus, in ACC, PV engagement is most consistently reduced in L2/3 and L5, with more prominent effects in females.

In the PrL, PV recruitment also decreased in layers 2/3 and 5, but the pattern depended on sex and subdivision (Fig. 3I-J; Table S2). In layer 2/3, males showed reduction after MDmc lesions, while females showed reductions after both MDmc and MDl compared with sham (Fig. 3J; Table S2). In layer 5, males again exhibited reduction after MDmc, whereas females showed robust reductions after both lesions (Fig. 3J; Table S2). Altogether, these data indicate that in PrL, MDmc, and, to a lesser extent, MDl lesions blunt PV interneuron recruitment by nociceptive stimulation, with effects that are strongest in layer 5 and more pronounced in females.

#### 4.3. SOM interneuron activation is selectively reduced in ACC layer 2/3 with minimal changes in PrL

We then assessed SOM interneuron activation using SOM⁺/cFos⁺ densities across layers in ACC and PrL (Fig. 4; Table S3).

In the ACC, SOM recruitment showed subdivision- and sex-dependent profiles (Fig. 4A-B). In layer 2/3, MDmc lesions reduced SOM⁺/cFos⁺ density in males compared with sham and MDl, whereas females showed no lesion-related differences (Table S3). In layer 5, females showed reduced SOM recruitment after both MDmc and MDl lesions compared with sham, while males showed little lesion effect (Fig. 4C; Table S3).

In the PrL (Fig. 4D-E), SOM interneurons show only limited changes in activation across layers, with no strong, layer-specific change of MDmc versus MDl lesions comparable to that observed in the ACC (Fig. 4F; Table S3). Overall, SOM engagement is most clearly disrupted in ACC, with comparatively limited modulation in PrL.

The excitotoxic lesions and immunohistochemical data indicate that MDmc and MDl shape prefrontal inhibitory microcircuits in distinct ways. To test their causal contribution to sensory and affective pain processing, we used projection-specific optogenetic inhibition of MD-ACC and MD-PrL pathways.

### 5. Validation of viral targeting and activity-dependent engagement of MD-PFC pathways

We carefully checked viral expression and optical fiber positioning to ensure the validity of optogenetic experiments (Fig. 5A, B, B1, C, C1). AAV5-CaMKII-driven opsin/GFP expression (Fig. 5B1, D-D1) was largely confined to the MD subdivisions (MDmc or MDl), with dense somatic labeling at the injection site and minimal spread to adjacent thalamic nuclei (Fig. 5B1, D1). Consistent with selective pathway targeting, cortical sections confirmed correct positioning of optical fibers above the intended ACC or PrL coordinates (Fig. 5C, E) and revealed a dense plexus of GFP-positive MD terminals within the illuminated territories (Fig. 5C1, E1). GFP-only illumination controls and distance-dependent cFos mapping are presented in Supplementary Figures S2-S4. At higher resolution, GFP-positive axons and boutons were observed surrounding cFos-positive neurons within the targeted PFC region (Fig. 5E2; Fig. S4), suggesting stimulation of local neuronal populations by MD terminal stimulation (Fig. S4A-H).

**Figure 5.**
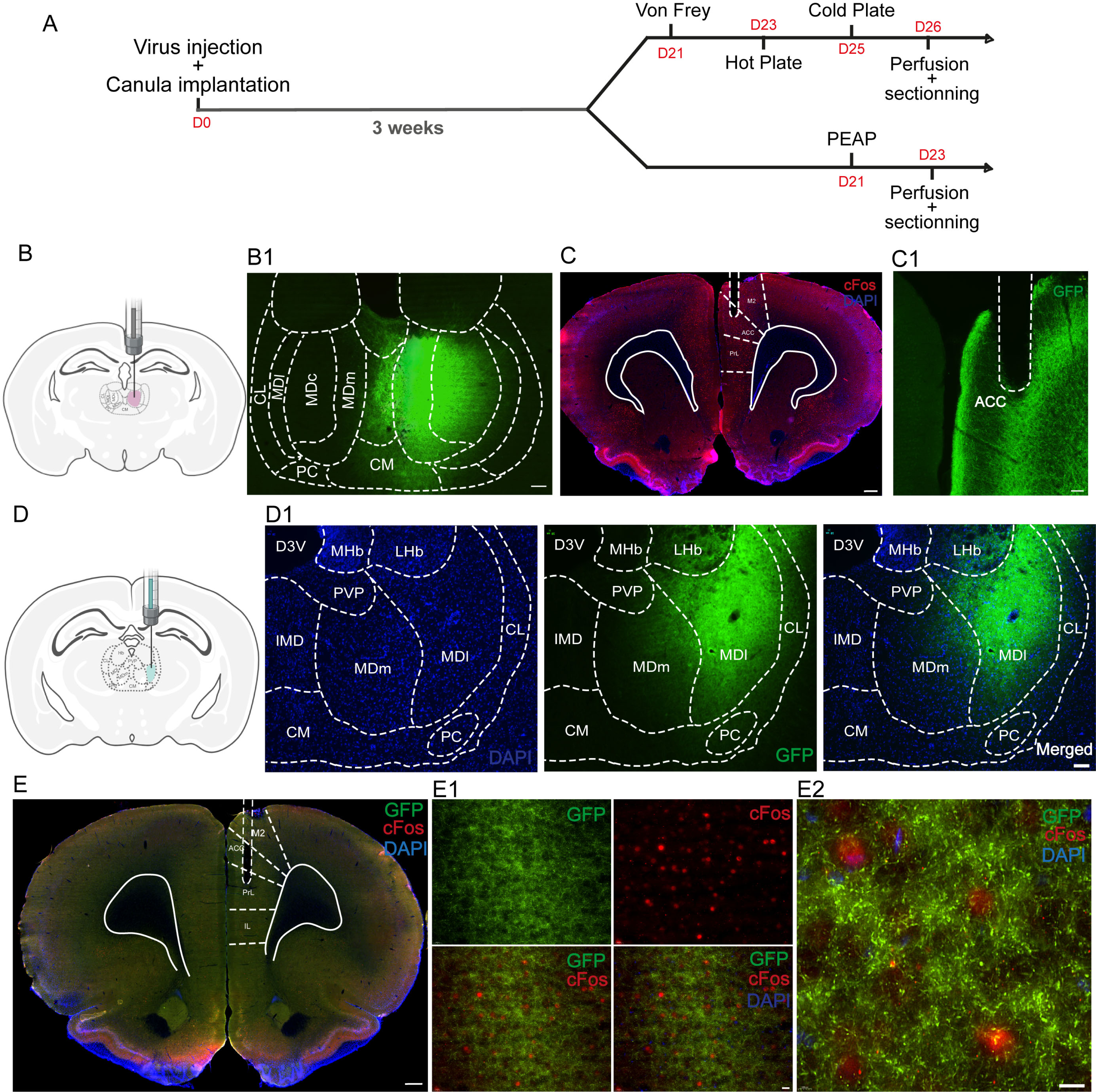
Optogenetic experimental design and viral targeting of MD subdivisions. **(A)** Timeline of the optogenetic protocol. **(B1)** Schematic of the stereotaxic viral injection into the MDmc. **(B2)** Coronal thalamic section showing the MDmc injection extent over atlas-based borders of MD subdivisions and neighboring nuclei. Scale bar: 70 μm. **(C1)** Low-magnification coronal section of the optical fiber placement site at ACC. DAPI (blue). cFos (red) Scale bar: 500 μm. **(C2)** Higher-magnification view of the fiber track and GFP⁺ MD terminals in ACC. Scale bar: 120 μm. **(D1)** Schematic coronal section shows the site of viral injection in the MDl. **(D2)** Coronal thalamic section showing the MDl injection site; Left (DAPI); Middle (GFP^+^); Right (Merged). Scale bar: 100 μm. **(E1)** Coronal section showing PrL implantation site and GFP⁺ MD terminals with cFos and DAPI. Scale bar: 500 μm. **(E2)** Enlarged PrL views showing GFP, cFos, and merged channels (±DAPI), illustrating the spatial overlap of MD terminals with activity-marked neurons. Scale bar: 20 μm. **(E3)** High-magnification confocal image showing GFP-positive axons and boutons coursing around and along cFos-expressing neuronal somata, with DAPI nuclear staining. Scale bar: 10 μm. Abbreviations: ACC, anterior cingulate cortex; PrL, prelimbic cortex; IL, infralimbic cortex; M2, secondary motor cortex; MD, mediodorsal thalamic nucleus; MDm, medial MD; MDl, lateral MD; MDc, central MD; IMD, intermediodorsal nucleus; CM, central medial nucleus; CL, central lateral nucleus; PC, paracentral nucleus; PVP, paraventricular posterior nucleus; MHb, medial habenula; LHb, lateral habenula; D3V, dorsal third ventricle.

### 6. Inhibition of MDmc and MDl projections to ACC unmasks latent hypersensitivity

We performed projection-specific optogenetic inhibition to transiently suppress MD terminal activity in prefrontal targets and directly test the causal contribution of this MD-PFC pathway to mechanical and thermal pain responses in males and females (Fig. 6).

**Figure 6.**
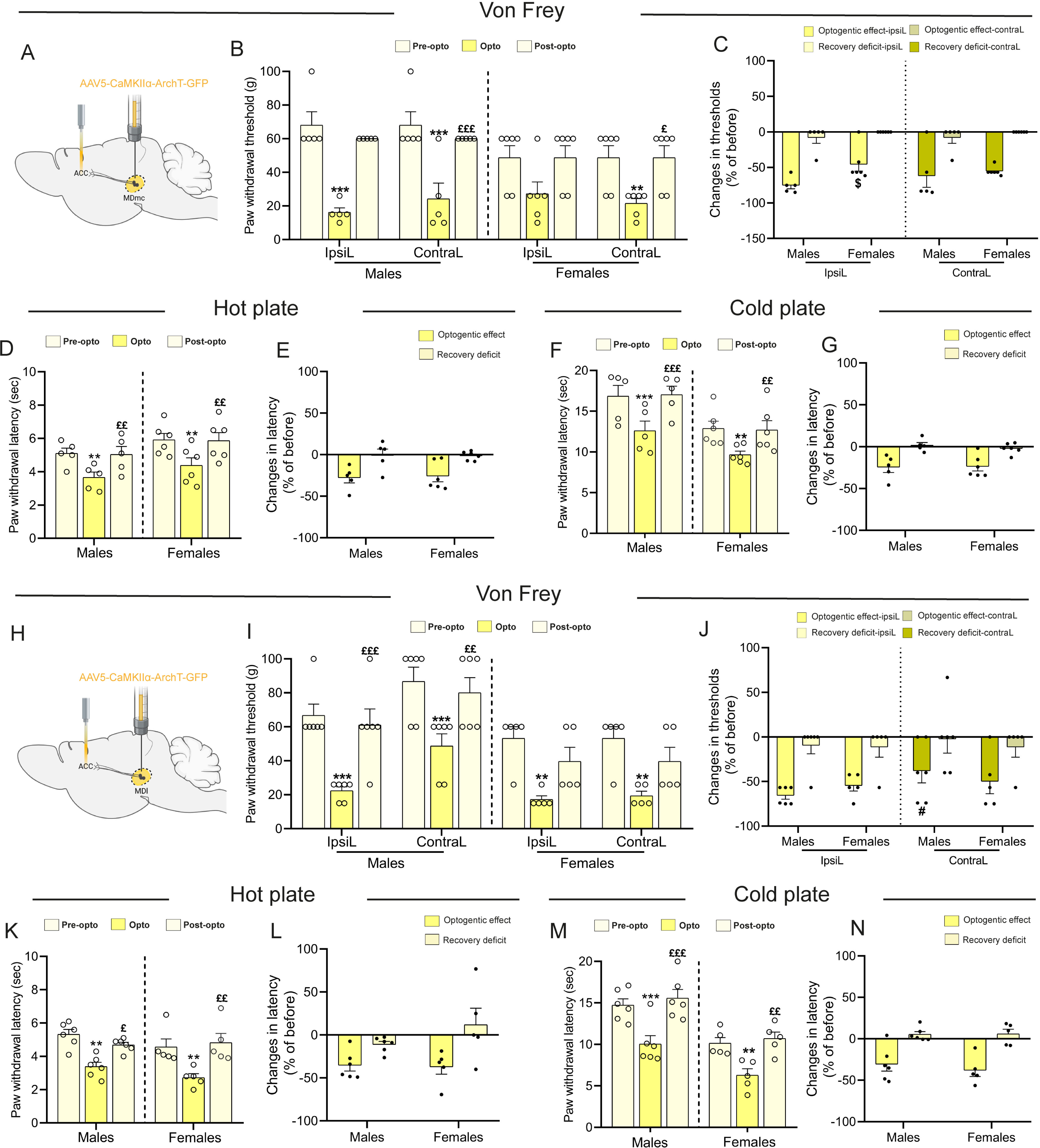
Optogenetic inhibition of MDmc and MDl-ACC projections exacerbates mechanical and thermal sensitivity. (A-G: MDmc-ACC cohort) **(A)** Schematic illustration of the experimental design showing AAV5-CaMKIIα-ArchT-GFP injection into the MDmc and optic-fiber implantation targeting its projection to the ACC. **(B)** Paw withdrawal thresholds (g) in the Von Frey test of the ipsilateral (IpsiL) and contralateral (ContraL) hind paws. **(C)** Percentage change in Von Frey thresholds relative to the individual pre-opto baseline for each sex and paw. **(D)** Paw withdrawal latencies (s) in the hot-plate test. **(E)** Percentage change in hot-plate withdrawal latency. **(F)** Paw withdrawal latencies (s) in the cold-plate test. **(G)** Percentage change in cold-plate withdrawal latency. (H-N, MDl-ACC cohort) **(H)** Schematic illustration of the experimental design showing AAV5-CaMKIIα-ArchT-GFP injection into the MDl and optic-fiber implantation targeting its projection to the ACC. **(I)** Paw withdrawal thresholds in the Von Frey test of the IpsiL and ContraL hind paws. **(J)** Percentage change in Von Frey thresholds relative to the individual pre-opto baseline for each sex and paw. **(K)** Paw withdrawal latencies in the hot-plate test. **(L)** Percentage change in hot-plate withdrawal latency. **(M)** Paw withdrawal latencies in the cold-plate test. **(N)** Percentage change in cold-plate withdrawal latency. Data are presented as mean ± SEM. MDmc-ACC-ArchT (males n = 5, females n = 6) and MDl-ACC-ArchT (males n = 6, females n = 5). Behavioral data were analyzed by RM two-way ANOVA (stage × sex) with Holm-Šidák *post-hoc* tests. During the stimulation epoch, von Frey normalized (%) responses were analyzed by two-way ANOVA (sex × paw side), and hot- and cold-plate measures were compared using unpaired t-tests (or Mann-Whitney when normality assumptions were not met). **p < 0.01, ***p < 0.001 Pre-opto vs Opto; £p < 0.05, ££p < 0.01, £££p < 0.001 Opto vs Post-opto; $p < 0.05 males vs females; #p < 0.05 ipsiL vs contraL paw during stimulation.

#### 6.1. Sensory-discriminative component: MDmc-ACC inhibition

##### a) Mechanical sensitivity

We first tested whether inhibiting the MDmc-ACC pathway alters von Frey responses (Fig. 6A). Two-way RM ANOVA analysis revealed a main effect of optogenetic condition (*F*_(5,44)_ = 19.56, p < 0.001) on von Frey withdrawal thresholds, and sex × optogenetic condition interaction (*F*_(5,44)_ = 3.72, p = 0.007), whereas the main effect of sex was not significant (*F*_(1,44)_ = 0.42, p = 0.534) (Fig. 6B). MDmc-ACC inhibition significantly decreased mechanical thresholds in males during optogenetic stimulation for both paws, ipsilateral (t = 6.46, p < 0.001) and contralateral (t = 6.46, p < 0.001) to the light delivery, compared to pre-stimulation baseline. After light offset, thresholds returned to baseline on the contralateral paw (t = 0.998, p = 0.691), whereas the ipsilateral paw remained different from baseline, indicating only partial recovery (t = 4.216, p < 0.001). A significant reduction in mechanical threshold was found also in females for the contralateral paw (t = 3.71, p = 0.009), that recovered after light offset (t = 0.0999, p = 1.000), whereas the ipsilateral comparison did not reach significance (t = 2.94, p = 0.056) (Fig. 6B). Baseline sex comparisons did not reach significance for either paw (ipsilateral: t = 1.923, p = 0.065; contralateral: t = 1.923, p = 0.065). We compared changes between ipsi- and contralateral paws and between males and females by normalizing the results against the pre-optogenetic baseline. The magnitude of the mechanical hypersensitivity did not differ as a function of paw side (*F*_(1,9)_ = 2.35, p = 0.159) or sex (*F*_(1,9)_ = 2.63, p = 0.140) (Fig. 6C). Overall, normalized data support a comparable reduction in mechanical thresholds across ipsilateral and contralateral paws, with no robust sex-dependent difference across conditions.

##### **b)** Heat sensitivity

We next asked whether similar effects would be observed for heat nociception (Fig. 6D and E). The ANOVA analysis revealed a main effect of optogenetic condition (*F*_(2,18)_ = 18.14, *p* < 0.001) on withdrawal latencies. In contrast, the main effect of sex (*F*_(1,18)_= 2.47, p = 0.151) and the interaction sex × optogenetic condition were not significant (*F*_(2,18)_ = 0.02, p = 0.980). A significant decrease in paw withdrawal latency was observed during optogenetic inhibition relative to the pre-stimulation baseline in males (t = 3.48, p = 0.008) and females (t = 4.06, p = 0.002). Importantly, after light offset, the post-stimulation phase did not differ from the pre-stimulation baseline in either sex (males: t = 0.15, p = 0.886; females: t = 0.13, p = 0.896), indicating a return to the baseline once illumination ended. Additionally, sex comparisons with normalized data did not reveal a difference between males and females (t(9) = -0.21, p = 0.837; Fig. 6E). Together, the hot plate data indicate that MDmc-ACC inhibition produces a robust but reversible facilitation of heat sensitivity, with comparable effects in males and females.

##### **c)** Cold sensitivity

Cold plate responses revealed a main effect of optogenetic condition (*F*_(2,18)_ = 22.01, p < 0.001) on withdrawal latencies, whereas the main effect of sex (*F*_(1,18)_ = 9.11, p = 0.015) and the interaction (*F*_(2,18)_ = 0.63, p = 0.544) did not reach significance (Fig. 6F). Optogenetic inhibition significantly reduced cold plate responses during stimulation relative to the pre-optogenetic phase in both males (t = 4.43, p < 0.001) and females (t = 3.67, p = 0.005). Importantly, responses returned to the baseline after stimulation offset in males (t = 0.18, p = 0.853) and females (t = 0.20, p = 0.837). Normalization against the pre-optogenetic baseline reveals that, changes did not differ between sexes (t(9) = -0.10, p = 0.921) (Fig. 6G). The above findings show that silencing MDmc inputs to ACC is sufficient to transiently enhance mechanical and thermal sensitivity.

#### 6.2. Sensory-discriminative component: MDl-ACC inhibition

##### a) Mechanical sensitivity

To assess how MDl-ACC inhibition affects mechanical sensitivity, paw withdrawal thresholds were measured following optogenetic inhibition (Fig. 6H). The analysis revealed a main effect of optogenetic condition (*F*_(5,45)_ = 17.84, p < 0.001) on withdrawal thresholds and a main effect of sex (*F*_(1,45)_= 16.32, p = 0.003), while the interaction did not reach significance (*F*_(5,45)_ = 2.21, p = 0.069). Optogenetic inhibition of MDl-ACC projections significantly lowered von Frey mechanical thresholds relative to the pre-opto phase in both sexes, affecting both paws (Fig. 6I). In males, thresholds decreased on the paw ipsilateral to light delivery (t = 5.19, p < 0.001) and the contralateral paw (t = 4.46, p < 0.001). Females showed the same pattern, with reductions on the ipsilateral (t = 3.85, p = 0.005) and contralateral paws (t = 3.85, p = 0.005). After light offset, thresholds returned toward baseline in both paws in both sexes, as the post-stimulation phase did not differ from the pre-stimulation baseline (males: ipsilateral t = 0.66, p = 0.510; contralateral t = 0.78, p = 0.685; females: ipsilateral t = 1.45, p = 0.631; contralateral t = 1.45, p = 0.485). Analysis of normalized data (Fig. 6J) did not reveal an effect of sex (*F*_(1,9)_= 0.001, p = 0.976), a paw effect (*F*_(1,9)_= 3.46, p = 0.096), or a sex × side interaction (*F*_(1,9)_= 1.72, p = 0.222). *Post-hoc* testing identified a significant ipsi-contra difference in males, indicating that mechanical sensitivity increased more on the contralateral paw than on the paw ipsilateral to light delivery (t = 2.35, p = 0.043). Overall, the von Frey data indicate that MDl-ACC, like MDmc-ACC, inhibition lowers mechanical thresholds reversibly.

##### **b)** Heat sensitivity

The analysis of hot plate latencies revealed a main effect of optogenetic condition (*F*_(2,18)_ = 15.32, p < 0.001) on withdrawal latencies, whereas the effect of sex (*F*_(1,18)_ = 3.56, p = 0.092) and the interaction (*F*_(2,18)_ = 0.86, p = 0.442) were not significant (Fig. 6K). *Post-hoc* tests indicated that optogenetic inhibition reduced paw withdrawal latency during stimulation relative to the pre-optogenetic phase in males (t = 3.82, p = 0.004) and females (t = 3.32, p = 0.008). Latencies returned to the baseline after light offset in both males (t = 1.25, p = 0.227) and females (t = 0.47, p = 0.645). The normalized hot plate effect indicated no differences between males and females (t(9) = 0.17, p = 0.870) (Fig. 6L). These findings indicate that MDl-ACC inhibition produces a reversible facilitation of heat sensitivity, similar to MDmc-ACC silencing.

##### **c)** Cold sensitivity

The cold plate test revealed a main effect of optogenetic condition (*F*_(2,18)_ = 23.60, p < 0.001) on withdrawal latencies and of sex (*F*_(1,18)_ = 25.57, p < 0.001), whereas the interaction between the two factors was not significant (*F*_(2,18)_ = 0.27, p = 0.764) (Fig. 6M). In males, optogenetic inhibition decreased cold plate responses during stimulation relative to baseline (t = 4.44, p < 0.001), and responses returned to baseline after light offset (t = 0.79, p = 0.441). In females, the same pattern was observed, with a reduction during stimulation (t = 3.33, p = 0.007) and recovery right after (t = 0.48, p = 0.635). Data normalization showed no differences between males and females (t(9) = 0.65, p = 0.534) (Fig. 6N).

Taken together, inhibition of MDmc-ACC and MDl-ACC projections demonstrates that both pathways are critically involved in shaping the sensory-discriminative component of pain. Transient silencing of either pathway acutely lowers mechanical and thermal thresholds in a reversible manner, indicating that ongoing MD-ACC activity is required to maintain normal nociceptive sensitivity across modalities and sexes.

### 7. MDl-ACC activation induces antinociception while MDmc-ACC drives hypersensitivity

We next asked whether enhancing, instead of silencing, MD output to ACC can dynamically tune pain processing in the opposite direction. For this purpose, we used ChR2-mediated specific activation of MDmc-ACC or MDl-ACC pathways in males and females.

#### 7.1. Sensory-discriminative component: MDmc-ACC activation

##### a) Mechanical sensitivity

To evaluate how selective activation of MDmc inputs to ACC (Fig. 7A) influences mechanical sensitivity, we applied von Frey stimulations. The ANOVA test (Fig. 7B) revealed a main effect of sex (*F*_(1,45)_= 11.11, p = 0.009) and optogenetic condition (*F*_(5,45)_= 4.22, p = 0.003) on withdrawal thresholds, with a significant sex × optogenetic condition interaction (*F*_(5,45)_ = 3.72, p = 0.007) (Fig. 7B). In males, the activation of the MDmc projections to the ACC reduced ipsilateral paw mechanical thresholds during stimulation relative to baseline (t = 3.39, p = 0.019) and recovered after light offset (t = 0, p = 1), whereas the contralateral paw change did not reach significance (t = 2.93, p = 0.056). In females, stimulation did not produce a reliable within-paw change on either side (ipsilateral: t = 1.91, p = 0.541; contralateral: t = 1.08, p = 0.904). The normalization of von Frey values against the pre-optogenetic condition (Fig. 7C) shows no differences between ipsi- and contralateral paws mechanical thresholds (*F*_(1,9)_ = 0.04, p = 0.844), nor between males and females (*F*_(1,9)_ = 0.03, p = 0.870).

**Figure 7.**
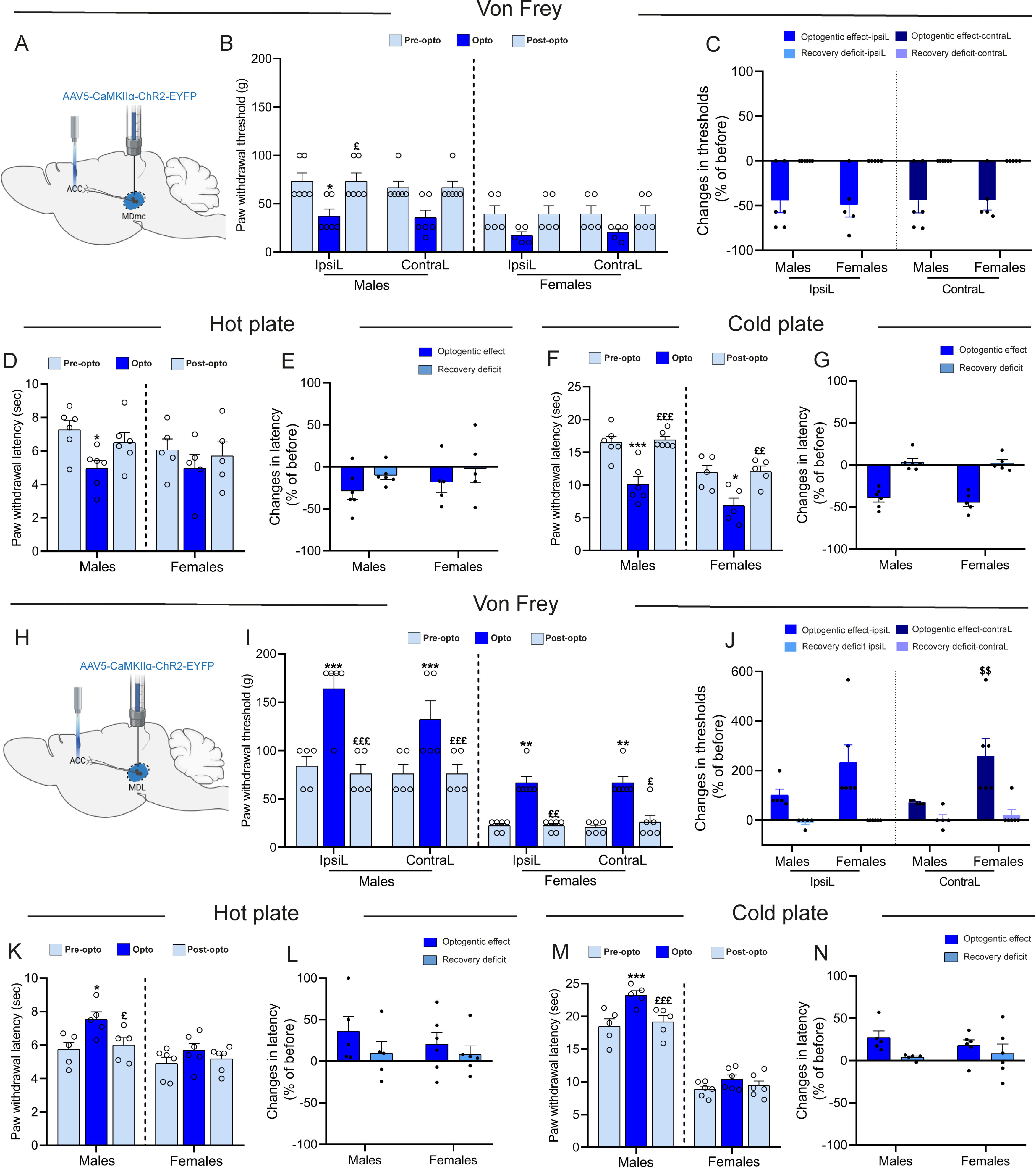
Optogenetic activation of MDmc and MDl-ACC projections differentially modulates mechanical and thermal sensitivity. (A-G, MDmc-ACC cohort) **(A)** Schematic illustration of the experimental design showing AAV5-CaMKIIα-ChR2-EYFP injection into the MDmc and optic-fiber implantation targeting its projection to the ACC. **(B)** Paw withdrawal thresholds (g) in the Von Frey test of the IpsiL and ContraL paws. **(C)** Percentage change in Von Frey thresholds relative to the individual pre-opto baseline. **(D)** Paw withdrawal latencies (s) in the hot-plate test. **(E)** Percentage change in hot-plate withdrawal latency (% of pre-opto). **(F)** Paw withdrawal latencies (s) in the cold-plate test. **(G)** Percentage change in cold-plate withdrawal latency (% of pre-opto). (H-N, MDl-ACC cohort) **(H)** Schematic illustration of the experimental design showing AAV5-CaMKIIα-ChR2-EYFP injection into the MDl and optic-fiber implantation targeting its projection to the ACC. **(I)** Paw withdrawal thresholds in the Von Frey test of the IpsiL and ContraL paws. **(J)** Percentage change in Von Frey thresholds relative to the individual pre-opto baseline. **(K)** Paw withdrawal latencies in the hot-plate test. **(L)** Percentage change in hot-plate withdrawal latency. **(M)** Paw withdrawal latencies in the cold-plate test. **(N)** Percentage change in cold-plate withdrawal latency. Data are presented as mean ± SEM. MDmc-ACC-ChR2 (males n = 6, females n = 5) and MDl-ACC-ChR2 (males n = 5, females n = 6). Behavioral data were analyzed by RM two-way ANOVA (stage × sex) with Holm-Šidák *post-hoc* tests. During the stimulation epoch, von Frey normalized (%) responses were analyzed by two-way ANOVA (sex × paw side), and hot- and cold-plate measures were compared using unpaired t-tests (or Mann-Whitney when normality assumptions were not met). *p < 0.05, **p < 0.01, ***p < 0.001 Pre-opto vs Opto; £p < 0.05, ££p < 0.01, £££p < 0.001 Opto vs Post-opto; $$p < 0.01 males vs females.

##### **b)** Heat sensitivity

Heat sensitivity was assessed by measuring hot-plate withdrawal latencies during MDmc-ACC activation. The analysis revealed a main effect of optogenetic condition (*F*_(2,18)_ = 4.78, p = 0.022). In contrast, the main effect of sex (*F*_(1,18)_ = 1.05, p = 0.333) and the interaction between the two factors (*F*_(2,18)_ = 0.63, p = 0.546) were not significant (Fig. 7D). In males, light delivery was accompanied by a decrease in withdrawal latency compared to the baseline (t = 3.06, p = 0.020) and recovered after light offset (t = 0.99, p = 0.331). In females, activation of MDmc-ACC projections did not affect heat latencies (t = 1.31, p = 0.499). We next compared light-induced changes between males and females, after normalization of withdrawal latencies against pre-optogenetic condition (Fig. 7E). Unpaired t-test showed no sex difference (t(9) = -0.68, p = 0.513), indicating a similar modulation across sexes.

##### **c)** Cold sensitivity

Cold-evoked response evaluated by the cold plate revealed a main effect of optogenetic condition (*F*_(2,17)_ = 26.41, p < 0.001) on withdrawal latencies and a main effect of sex (*F*_(1,17)_= 9.51, p = 0.010), whereas interaction was not significant (*F*_(2,17)_ = 1.17, p = 0.333) (Fig. 7F). In males, illumination lowered the cold plate latencies compared to the baseline value (t = 5.29, p < 0.001), that recovered after light offset (t = 0.33, p = 0.745). In females, a similar stimulation-induced decrease was observed (t = 3.14, p = 0.012), followed by a total recovery (t = 1.28, p = 0.218). Normalized light-evoked changes during stimulation did not differ between males and females (t(9) = 0.85, p = 0.416), indicating that the magnitude of the light-driven modulation was comparable across sexes (Fig. 7G).

Taken together, von Frey, hot plate, and cold plate data indicate that MDmc-ACC activation consistently shifts responses in the same direction as ArchT-mediated inhibition. Thus, for MDmc-ACC, both decreasing and increasing pathway activity shift nociceptive thresholds toward hypersensitivity.

#### 7.2. Sensory-discriminative component: MDl-ACC activation

##### a) Mechanical sensitivity

Optogenetic activation of the MDl-ACC pathway (Fig. 7H) resulted in a main effect of sex (*F*_(1,44)_ = 84.72, p < 0.001) and a main effect of optogenetic condition (*F*_(5,44)_ = 24.77, p < 0.001) on mechanical withdrawal thresholds (Fig. 7I), while the interaction did not reach significance (*F*_(5,44)_ = 2.28, p = 0.063) (Fig. 7I). *Post-hoc* tests revealed that, in males, thresholds increased during stimulation relative to baseline for both ipsilateral (t = 6.28, p < 0.001) and contralateral (t = 4.40, p < 0.001) paws, and recovered after light offset. In females, the activation of MDl-ACC projections similarly elevated thresholds during stimulation in the ipsilateral (t = 3.81, p = 0.006) and the contralateral (t = 3.97, p = 0.004) paw. Normalization of von Frey responses against the baseline (Fig. 7J) did not show a significant effect of sex (*F*_(1,9)_ = 4.16, p = 0.072), paw side (*F*_(1,9)_ = 0.00, p = 0.978), or a sex × paw side interaction (*F*_(1,9)_= 2.15, p = 0.177).

##### **b)** Heat sensitivity

Hot plate testing revealed a main effect of sex (*F*_(1,18)_ = 10.63, p = 0.010) and a main effect of optogenetic condition (*F*_(2,18)_ = 6.72, p = 0.007) on withdrawal latencies, whereas the sex × condition interaction was not significant (*F*_(2,18)_ = 1.25, p = 0.310) (Fig. 7K). Further analysis in males showed that the activation of MDl-ACC projections increased withdrawal latencies compared to baseline (t = 3.27, p = 0.013), that returned to baseline after light offset (t = 0.47, p = 0.642). In females, we did not detect any significant effect of optogenetic stimulation (t = 1.56, p = 0.355). Normalization against the baseline (Fig. 7L) indicates that males and females exhibited a similar amplitude of change during light stimulation (t(9) = 0.70, p = 0.504).

##### **c)** Cold sensitivity

The analysis of cold-plate responses revealed main effects of sex (*F*_(1,18)_ = 156.12, p < 0.001) and optogenetic condition (*F*_(2,18)_ = 15.62, p < 0.001) on withdrawal latencies, along with a significant sex × phase interaction (*F*_(2,18)_ = 4.64, p = 0.024) (Fig. 7M). In males, the cold-plate measure was increased during light stimulation compared with baseline (t = 5.38, p < 0.001), that recovered right after stimulation ended (t = 0.75, p = 0.463). In females, there was no significant effect induced by light stimulation (t = 1.91, p = 0.202). Normalization of light-evoked changes showed comparable effects of optogenetic stimulation in both sexes (t(9) = 0.92, p = 0.383) (Fig. 7N) Activation of MDl-ACC projections produced a sensory pattern opposite to inhibition, with higher mechanical thresholds and longer heat latencies during light delivery.

### 8. Inhibition of MD-PrL outputs exacerbates sensory nociception

Because lesions, tracing, and cFos/PV/SOM mapping all pointed to partially segregated MDmc and MDl outputs onto ACC and PrL, we next asked whether the causal influence of MD-prefrontal drive on pain processing was restricted to ACC or extended to PrL as a parallel control node. To address this, we applied the same ArchT-based inhibition strategy to MDmc-PrL and MDl-PrL pathways and examined its impact on sensory-discriminative nociception.

#### 8.1. Sensory-discriminative component: MDmc-PrL inhibition

##### a) Mechanical sensitivity

Mechanical sensitivity was assessed across optogenetic epochs and sex (Fig. 8A). The ANOVA revealed a main effect of sex (*F*_(1,50)_ = 15.569, p = 0.003) and a main effect of optogenetic condition (*F*_(5,50)_ = 12.435, p < 0.001) on withdrawal thresholds, with no interaction between the two factors (*F*_(5,50)_ = 0.549, p = 0.738) (Fig. 8B). *Post-hoc* analyses showed that in males, optogenetic stimulation induced a clear, reversible reduction in mechanical thresholds on both ipsilateral paw (t = 4.53, p < 0.001) and the contralateral paw (t = 3.71, p = 0.006) and returned to baseline after light offset (ipsilateral: t = 1.30, p = 0.022; contralateral: t = 0, p = 1). In females, optogenetic modulation was restricted to the ipsilateral paw, with a significant reduction during light stimulation compared to baseline (t = 3.47, p = 0.015), reversible after light offset (t = 0.79, p = 0.966). The percent-normalized data indicate that the magnitude of change in mechanical sensitivity was comparable between ipsilateral and contralateral paws and did not differ between males and females. The ANOVA results confirmed that no effect of paw side (*F*_(1,10)_ = 0.00, p = 0.998), sex (*F*_(1,10)_ = 0.96, p = 0.351), or sex × paw side interaction (*F*_(1,10)_ = 0.71, p = 0.418) were detected (Fig. 8C).

**Figure 8.**
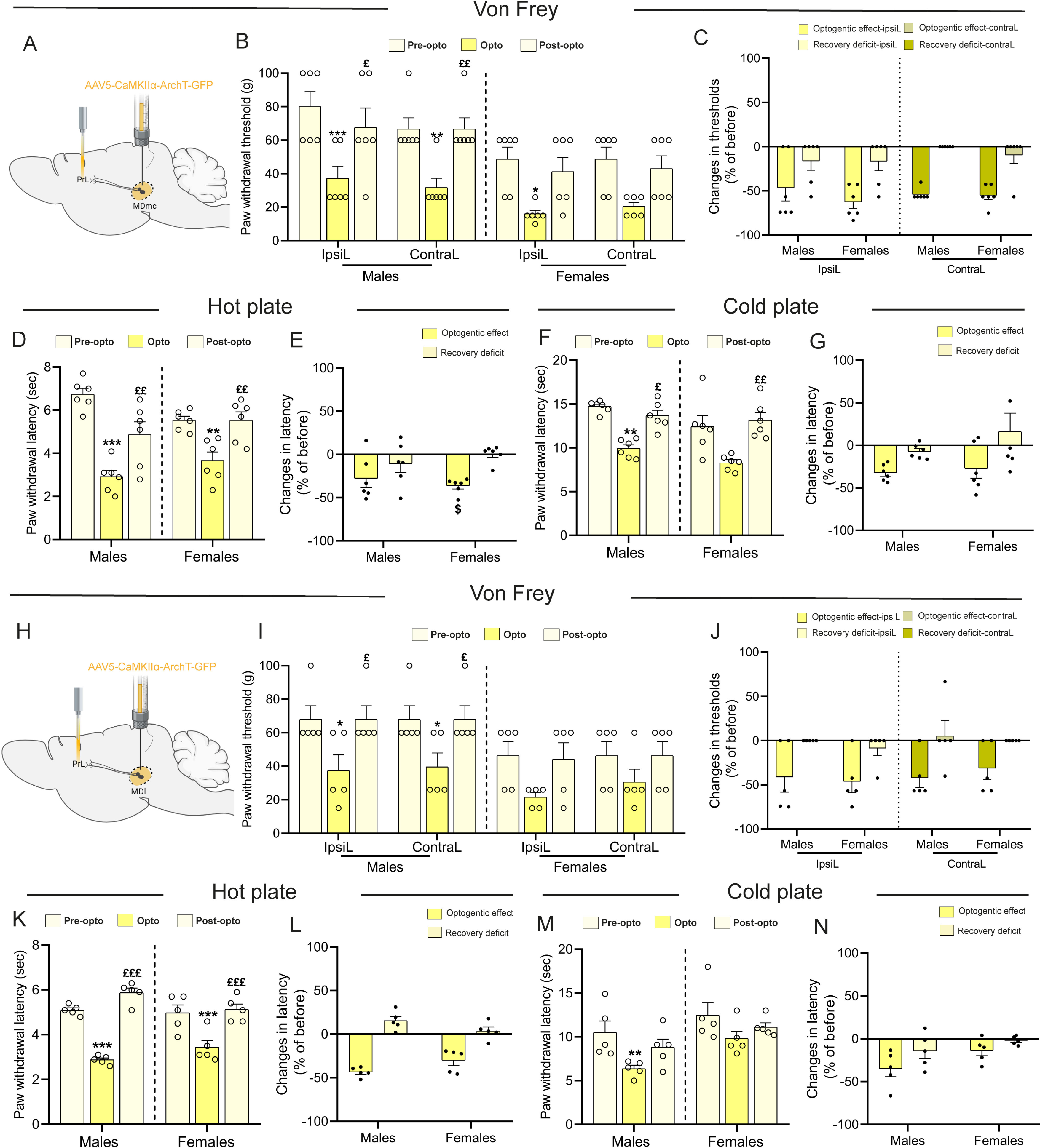
Optogenetic inhibition of MDmc and MDl-PrL projections lowers mechanical and thermal pain thresholds. (A-G, MDmc-PrL cohort) **(A)** Schematic illustration of the experimental design showing AAV5-CaMKIIα-ArchT-GFP injection into the MDmc and optic-fiber implantation targeting its projection to the PrL. **(B)** Paw withdrawal thresholds (g) in the Von Frey test of the IpsiL and ContraL paws. **(C)** Percentage change in Von Frey thresholds relative to the individual pre-opto baseline. **(D)** Paw withdrawal latencies (s) in the hot-plate test. **(E)** Percentage change in hot-plate withdrawal latency (% of pre-opto). **(F)** Paw withdrawal latencies (s) in the cold-plate test. **(G)** Percentage change in cold-plate withdrawal latency (% of pre-opto). (H-N, MDl-PrL cohort) **(H)** Schematic illustration of the experimental design showing AAV5-CaMKIIα-ArchT-GFP injection into the MDl and optic-fiber implantation targeting its projection to the PrL. **(I)** Paw withdrawal thresholds in the Von Frey test of the IpsiL and ContraL paws. **(J)** Percentage change in Von Frey thresholds relative to the individual pre-opto baseline. **(K)** Paw withdrawal latencies in the hot-plate test. **(L)** Percentage change in hot-plate withdrawal latency. **(M)** Paw withdrawal latencies in the cold-plate test. **(N)** Percentage change in cold-plate withdrawal latency. Data are presented as mean ± SEM. MDmc-PrL-ArchT (males n = 6, females n = 6) and MDl-PrL-ArchT (males n = 5, females n = 5). Behavioral data were analyzed by RM two-way ANOVA (stage × sex) with Holm-Šidák *post-hoc* tests. During the stimulation epoch, von Frey normalized (%) responses were analyzed by two-way ANOVA (sex × paw side), and hot- and cold-plate measures were compared using unpaired t-tests (or Mann-Whitney when normality assumptions were not met). *p < 0.05, **p < 0.01, ***p < 0.001 Pre-opto vs Opto; £p < 0.05, ££p < 0.01, £££p < 0.001 Opto vs Post-opto; $p < 0.05 males vs females.

##### **b)** Heat sensitivity

Hot-plate withdrawal latencies were next assessed. The analysis revealed an effect of optogenetic condition (*F*_(2,20)_ = 36.14, p < 0.001) on withdrawal latencies with a significant optogenetic condition × sex interaction (*F*_(2,20)_ = 5.23, p = 0.015), while the overall effect of sex was not significant (*F*_(1,20)_ = 0.04, p = 0.848) (Fig. 8D). In males, optogenetic inhibition reduced withdrawal latency during light delivery relative to baseline (t = 7.92, p < 0.001) and the recovery was partial after light offset (t = 3.87, p < 0.001). Females showed the same pattern, with reduced latency during stimulation (t = 3.87, p = 0.003) and a total recovery after light offset (t = 0.00, p = 1.000). The percent-normalized reduction in withdrawal latency was significantly greater in males than in females (t(10) = -2.79, p = 0.019), indicating a stronger facilitation of heat sensitivity during MDmc-PrL inhibition in males (Fig. 8E).

##### **c)** Cold sensitivity

During the cold plate assay, the ANOVA results revealed a main effect of sex (*F*_(1,20)_ = 8.60, p = 0.015) and optogenetic condition (*F*_(2,20)_ = 16.21, p < 0.001) on withdrawal latencies, with no interaction between the two factors (*F*_(2,20)_ = 0.44, p = 0.653) (Fig. 8F). Consistently, multiple comparisons indicated that, in males, withdrawal latency dropped during light delivery relative to the pre-light baseline (t = 3.78, p = 0.004) and recovered when the light turned off (t = 0.83, p = 0.416). The same pattern was observed in females, where latency decreased during photoinhibition compared with baseline (t = 3.37, p = 0.006) and recovered right after (t = 0.46, p = 0.650). Along the same line, the baseline-normalized light-evoked changes during stimulation were also comparable between males and females (t(10) = -0.40, p = 0.695) (Fig. 8G).

Taken together, ArchT-mediated inhibition of MDmc-PrL projections consistently lowers mechanical thresholds and hot- and cold-plate latencies in both sexes.

#### 8.2. Sensory-discriminative component: MDl-PrL inhibition

##### a) Mechanical sensitivity

The analysis of mechanical thresholds during optogenetic inhibition of MDl-PrL projections (Fig. 8H) revealed a main effect of optogenetic condition (*F*_(5,40)_ = 7.41, p < 0.001) and of sex (*F*_(1,40)_ = 6.32, p = 0.036), while the interaction was not significant (*F*_(5,40)_ = 0.34, p = 0.883) (Fig. 8I). In males, optogenetic inhibition induced a transient reduction in withdrawal thresholds during light delivery on both the ipsilateral (t = 3.24, p = 0.031) and contralateral paws (t = 3.01, p = 0.044), with thresholds returning toward baseline after light offset (ipsilateral: t = 3.24, p = 0.035; contralateral: t = 3.01, p = 0.049). In females, comparable within-paw modulations did not reach significance, indicating a weaker effect. The normalized MDl-PrL-ArchT data did not reveal any effect of sex (*F*_(1,8)_= 0.03, p = 0.861), nor paw side (*F*_(1,8)_= 0.46, p = 0.515), and the sex × paw side interaction was not significant (*F*_(1,8)_= 0.58, p = 0.467) (Fig. 8J), indicating a comparable modulation of mechanical sensitivity between males and females and between ipsilateral and contralateral paws during MDl-PrL inhibition.

##### **b)** Heat sensitivity

Heat withdrawal latency analysis revealed a main effect of optogenetic condition (*F*_(2,16)_ = 77.27, p < 0.001) on withdrawal latencies, no overall main effect of sex (*F*_(1,16)_ = 0.19, p = 0.673), and a significant sex × optogenetic condition interaction (*F*_(2,16)_ = 5.48, p = 0.015) (Fig. 8K). In males, ArchT-mediated inhibition reduced withdrawal latency during light delivery relative to baseline (t = 7.87, p < 0.001) and the recovery was partial (t = 2.77, p = 0.014). Females showed a similar drop during stimulation (t = 5.46, p < 0.001) and recovery (t = 0.50, p = 0.626). Normalized light-evoked change analysis showed a broadly comparable light-evoked change across sexes (t(8) = -2.08, p = 0.071; Fig. 8L).

##### **c)** Cold sensitivity

Withdrawal latencies in the cold plate test revealed a main effect of optogenetic condition (*F*_(2,16)_ = 8.43, p = 0.003) and sex (*F*_(1,16)_ = 7.33, p = 0.027) with no sex × condition interaction (*F*_(2,16)_ = 1.72, p = 0.210) (Fig. 8M). In males, photoinhibition lowered withdrawal latency relative to the pre-stimulation period (t = 4.16, p = 0.002) and latency recovered after light offset (t = 1.73, p = 0.103). In females, the photoinhibition did not affect withdrawal latency (t = 1.55, p = 0.366). Moreover, baseline-normalized light-evoked change did not differ between males and females (t(8) = -1.90, p = 0.094; Fig. 8N).

Overall, silencing MDl-PrL projections also produces a decrease in mechanical and thermal withdrawal latencies, with stronger and more consistent effects in males.

### 9. MDmc-PrL activation drives sensory hypersensitivity, while MDl supports analgesia

Next, we asked how driving the activation of MDmc and MDl projections to PrL would reshape pain processing.

#### 9.1. Sensory-discriminative component: MDmc-PrL activation

##### a) Mechanical sensitivity

Since both MDmc lesions and MDmc-PrL inhibition facilitated nocifensive behavior, we tested whether activating MDmc terminals in PrL would symmetrically dampen pain sensitivity (Fig. 9A). The analysis of von Frey responses revealed a main effect of optogenetic condition (*F*_(5,50)_ = 9.54, p < 0.001) and a main effect of sex (*F*_(1,50)_ = 6.13, p = 0.033) on withdrawal thresholds, with no interaction (*F*_(5,50)_ = 0.51, p = 0.765) (Fig. 9B). *Post-hoc* analyses indicated that in males, light delivery significantly reduced mechanical responsiveness on the ipsilateral paw relative to baseline (t = 4.16, p = 0.002), and returned to baseline after light offset (t = 1.10, p = 0.724). On the contralateral paw, withdrawal values also decreased during stimulation compared with baseline (t = 3.86, p = 0.004), and recovered after (t = 2.11, p = 0.278). In females, light delivery was associated with lower withdrawal values on both paws, but these changes did not reach significance (ipsilateral: t = 2.74, p = 0.112; contralateral: t = 2.01, p = 0.334). In the percent-normalized analysis, ANOVA analysis did not reveal a main effect of paw side (*F*_(1,50)_ = 0.29, p = 0.601), sex (*F*_(1,10)_ = 0.03, p = 0.867), nor the interaction (*F*_(1,50)_ = 0.97, p = 0.347) (Fig. 9C). Thus, the magnitude of the mechanical modulation remained comparable between ipsilateral and contralateral paws and did not differ between males and females.

**Figure 9.**
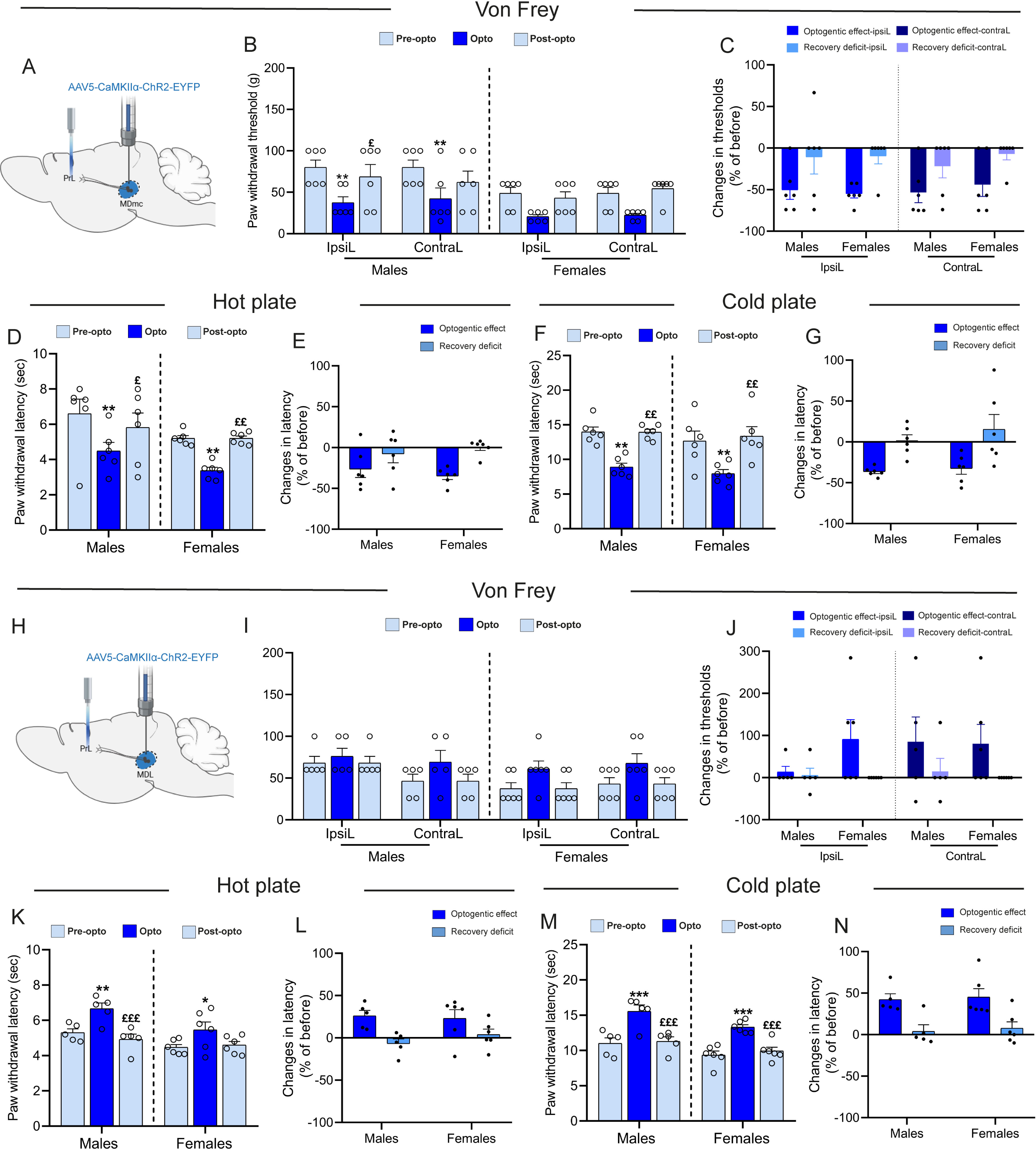
Optogenetic activation of MDmc-PrL projections enhances mechanical and thermal hypersensitivity, while MDl-PrL activation exerts inverse modulatory effects on nociceptive thresholds. (A-G, MDmc-PrL cohort) **(A)** Schematic illustration of the experimental design showing AAV5-CaMKIIα-ChR2-EYFP injection into the MDmc and optic-fiber implantation targeting its projection to the PrL. **(B)** Paw withdrawal thresholds (g) in the Von Frey test of the IpsiL and ContraL paws. **(C)** Percentage change in Von Frey thresholds relative to the individual pre-opto baseline. **(D)** Paw withdrawal latencies (s) in the hot-plate test. **(E)** Percentage change in hot-plate withdrawal latency (% of pre-opto). **(F)** Paw withdrawal latencies (s) in the cold-plate test. **(G)** Percentage change in cold-plate withdrawal latency (% of pre-opto). (H-N, MDl-PrL cohort) **(H)** Schematic illustration of the experimental design showing AAV5-CaMKIIα-ChR2-EYFP injection into the MDl and optic-fiber implantation targeting its projection to the PrL. **(I)** Paw withdrawal thresholds in the Von Frey test of the IpsiL and ContraL paws. **(J)** Percentage change in Von Frey thresholds relative to the individual pre-opto baseline. **(K)** Paw withdrawal latencies in the hot-plate test. **(L)** Percentage change in hot-plate withdrawal latency. **(M)** Paw withdrawal latencies in the cold-plate test. **(N)** Percentage change in cold-plate withdrawal latency. Data are presented as mean ± SEM. MDmc-PrL-ChR2 (males n = 6, females n = 6) and MDl-PrL-ChR2 (males n = 5, females n = 6). Behavioral data were analyzed by RM two-way ANOVA (stage × sex) with Holm-Šidák *post-hoc* tests. During the stimulation epoch, von Frey normalized (%) responses were analyzed by two-way ANOVA (sex × paw side), and hot- and cold-plate measures were compared using unpaired t-tests (or Mann-Whitney when normality assumptions were not met). *p < 0.05, **p < 0.01, ***p < 0.001 Pre-opto vs Opto; £p < 0.05, ££p < 0.01, £££p < 0.001 Opto vs Post-opto.

##### **b)** Heat sensitivity

Heat sensitivity analysis revealed a main effect of optogenetic condition (*F*_(2,20)_ = 16.51, p < 0.001) on withdrawal latencies, with no main effect of sex (*F*_(1,20)_ = 2.76, p= 0.127) or interaction (*F*_(2,20)_ = 0.59, p = 0.562) (Fig. 9D). In males, multiple comparison showed that blue-light delivery lowered withdrawal latency relative to the pre-stimulation period (t = 4.11, p = 0.002). Latency returned to baseline values after light offset (t = 1.52, p = 0.144). Females showed the same pattern, stimulation reduced withdrawal latency relative to pre-stimulation (t = 3.56, p = 0.006). Latency returned to baseline values after light offset (t = 0.00, p = 1.000). Furthermore, the percent-normalized modulation indicated that the magnitude of change did not differ between males and females (t(10) = 0.76, p = 0.463; Fig. 9E).

##### **c)** Cold sensitivity

Cold-plate withdrawal latencies analysis revealed a main effect of optogenetic condition on withdrawal latencies (*F*_(2,20)_ = 21.27, p < 0.001), with no main effect of sex (*F*_(1,20)_ = 1.27, p = 0.285) or interaction (*F*_(2,20)_ = 0.14, p = 0.871) (Fig. 9F). In males, blue-light delivery shortened cold-plate withdrawal latency relative to the pre-stimulation condition (t = 4.04, p = 0.002). Latency returned to baseline values after light offset (t = 0.03, p = 0.979). Females showed the same pattern, with a significant reduction in withdrawal latency during light delivery relative to pre-stimulation (t = 3.59, p = 0.004) and total recovery after light offset (t = 0.72, p = 0.483). During stimulation, males and females did not differ in the normalized light-evoked change (t(10) = -0.52, p = 0.615), indicating a comparable modulation magnitude across sexes (Fig. 9G).

Taken together, ChR2 activation of MDmc-PrL projections consistently shortens mechanical, heat- and cold-evoked withdrawal latencies in both sexes. Notably, this sensory pattern mirrors the hyperalgesia observed during MDmc-PrL inhibition.

#### 9.2. Sensory-discriminative component: MDl-PrL activation

We then turned to the lateral MD-PrL pathway, for which inhibition hyperalgesia, especially for thermal sensitivity (Fig. 9H).

##### **a)** Mechanical sensitivity

Mechanical withdrawal analysis revealed a main effect of optogenetic condition (*F*_(5,44)_ = 2.95, p = 0.022) and sex (*F*_(1,44)_ = 6.54, p = 0.031) on withdrawal thresholds, while the interaction was not significant (*F*_(5,44)_ = 1.09, p = 0.381) (Fig. 9I). The activation of MDl-PrL projections did not affect withdrawal values relative to the pre-stimulation period on either paw of both males (ipsilateral: t = 0.60, p = 0.996; contralateral: t = 1.71, p = 0.722) and females (ipsilateral: t = 1.95, p = 0.449; contralateral: t = 2.03, p = 0.449) (Fig. 9J).

##### **b)** Heat sensitivity

The analysis of hot-plate withdrawal latencies revealed a main effect of sex (*F*_(1,18)_ = 8.01, p = 0.020) and optogenetic condition (*F*_(2,18)_ = 14.12, p < 0.001) on withdrawal latencies, while the interaction was not significant (*F*_(2,18)_ = 1.31, p = 0.293) (Fig. 9K). In males, withdrawal latency increased during light delivery relative to the pre-light period (t = 3.40, p = 0.006) and returned to baseline after stimulation ended (t = 0.95, p = 0.355). Females showed the same pattern, with latencies increasing during light delivery compared to the pre-light period (t = 2.74, p = 0.040), and recovering after light offset (t = 0.37, p = 0.719). In addition, the analysis of the light-evoked, baseline-normalized change in hot-plate latency during stimulation showed no sex difference (Mann-Whitney U = 15.00, p = 1.000; Fig. 9L).

##### **c)** Cold sensitivity

Cold-plate withdrawal latencies analysis showed a main effect of optogenetic condition (*F*_(2,18)_ = 40.58, p < 0.001) and sex (*F*_(1,18)_ = 6.86, p = 0.028) on withdrawal latencies, with no interaction (*F*_(2,18)_ = 0.35, p = 0.706) (Fig. 9M). In males, blue-light delivery increased cold-evoked withdrawal latency relative to the pre-stimulation period (t = 5.91, p < 0.001) and values returned to baseline after light offset (t = 0.36, p = 0.720). A comparable pattern was observed in females, with higher latencies during light delivery (t = 5.65, p < 0.001) and total recovery after light offset (t = 0.81, p = 0.430). Baseline-normalized light-evoked change likewise did not differ between sexes (U = 10.00, p = 0.429), supporting a comparable magnitude of modulation in males and females (Fig. 9N).

Overall, in contrast to MDmc-PrL, where both inhibition and activation converge on hyperresponsiveness, MDl-PrL behaves more like a modality-sensitive regulator with inhibition favouring hyperalgesia and activation preferentially supporting analgesia for thermal inputs, with limited impact on mechanical sensitivity.

### 10. MDmc and MDl projections to ACC and PrL differentially modulate the affective-motivational component of pain

Since pain processing also involves an affective-motivational component, and because chronic MDmc and MDl lesions differentially affected escape/avoidance behavior in the PEAP task, we next investigated how MD subdivisions projecting to the mPFC contribute to this dimension, by comparing independent groups under light-off and light-on conditions.

#### 10.1. MD-ACC inhibition

To evaluate the effect of MDmc-ACC and MDl-ACC inhibition (Fig. 10A) on the escape/avoidance behavior, we evaluated time spent in the lit compartment using a three-way ANOVA and the analysis revealed a main effect of optogenetic condition (*F*_(1,32)_ = 13.88, p < 0.001) and a main effect of subdivision (*F*_(1,32)_ = 26.44, p < 0.001), whereas the main effect of sex did not reach significance (*F*_(1,32)_ = 2.93, p = 0.100). Importantly, the only significant interaction was between the optogenetic condition and the subdivision (*F*_(1,32)_ = 19.35, p < 0.001) (Fig. 10B). In males of the MDmc group, light-induced inhibition amplified PEAP responses (t = 3.70, p = 0.020), whereas they remained unchanged in the MDl group (t = 0.57, p > 0.990). The same pattern was observed in females, where inhibition in the ACC increases escape/avoidance behavior in the MDmc (t = 4.42, p = 0.002) but not MDl (t = 1.25, p = 0.960) group. These results are consistent with reduced aversion/avoidance engagement under MDl-ACC inhibition even in the context of heightened nociceptive sensitivity.

**Figure 10.**
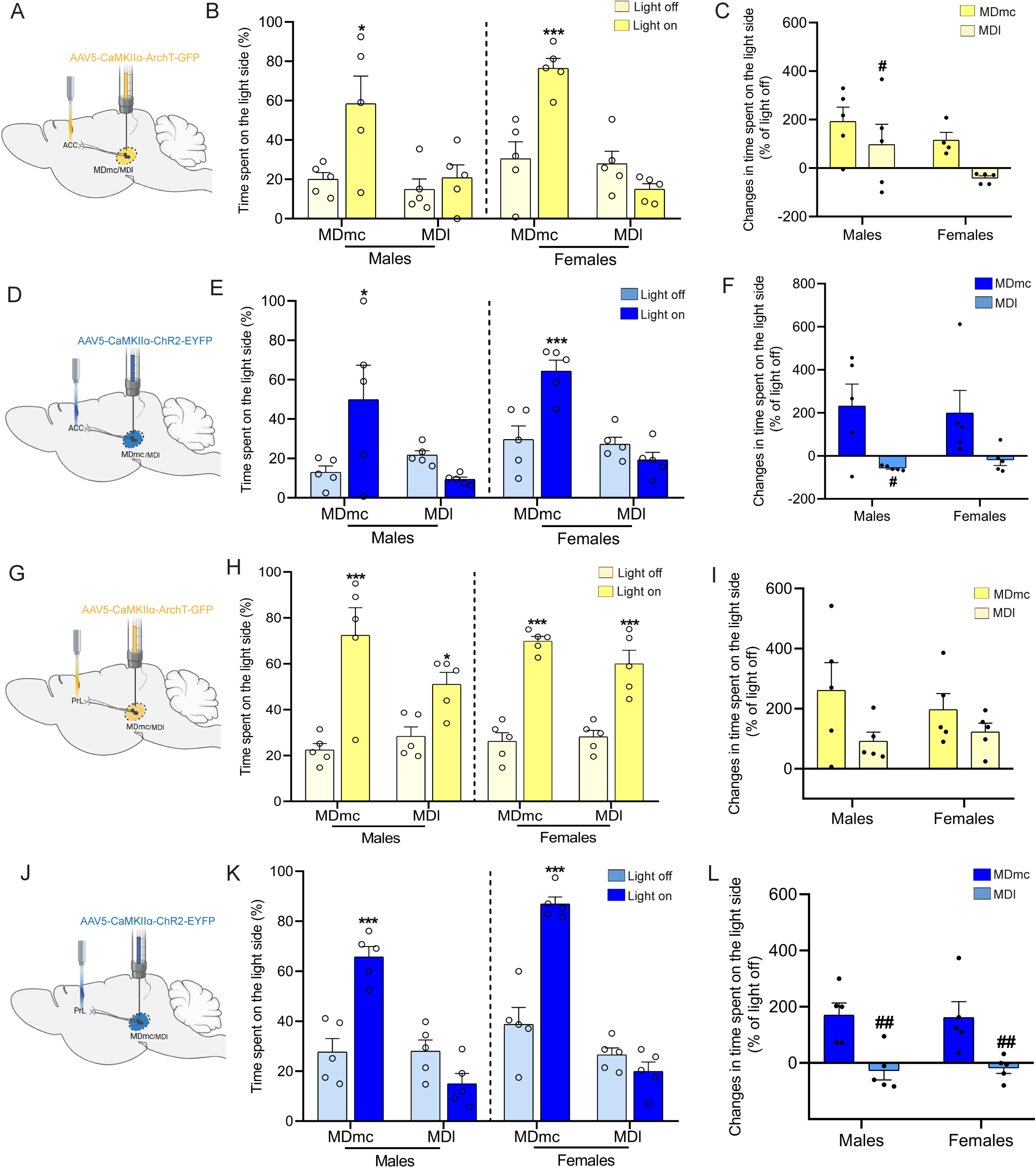
Optogenetic inhibition and activation of MD projections to ACC and PrL uncover opposing contributions of MD-PFC pathways to affective pain processing. ACC manipulations **(A)** Schematic illustration of the inhibitory experiment showing AAV5-CaMKIIα-ArchT-GFP injection into the MD (MDmc or MDl) and optic-fiber implantation over the ACC. **(B)** Time spent on the light side (%) for MDmc and MDl groups under light-off and light-on conditions. **(C)** Percentage change in time spent on the light side relative to the individual light-off baseline. **(D)** Schematic illustration of the activation experiment showing AAV5-CaMKIIα-ChR2-EYFP injection into MDmc or MDl and optic-fiber implantation over the ACC. **(E)** Time spent on the light side for MDmc and MDl groups with light off and light on during ChR2 activation. **(F)** Percentage change in time on the light side (% of light-off baseline). PrL manipulations **(G)** Schematic of the inhibitory experiment with AAV5-CaMKIIα-ArchT-GFP injection into MDmc or MDl and optic-fiber implantation over the PrL. **(H)** Time spent on the light side (%) for MDmc and MDl groups under light-off and light-on conditions during MD-PrL inhibition. **(I)** Percentage change in time spent on the light side (% of light-off baseline. **(K)** Time spent on the light side (%) for MDmc and MDl groups under light-off and light-on conditions during MD-PrL activation. **(L)** Percentage change in time on the light side (% of light-off baseline). Data are presented as mean ± SEM. n: 5 per group. Time spent on the light side was analyzed using a three-way ANOVA with factors subdivision, optogenetic condition, and sex. Percentage change in time on the light side was analyzed using a two-way ANOVA with factors subdivision and sex; significant effects were explored with Holm-Šidák *post-hoc* multiple-comparisons tests. *p < 0.05, ***p < 0.001 light on vs light off within the same subdivision; #p < 0.05, ##p < 0.01, ###p < 0.001 MDl vs MDmc under the same optogenetic condition.

We next normalized the PEAP responses against the light-off condition for each subdivision and sex, to compare changes in light-induced modulation between MDmc and MDl. The analysis revealed a main effect of subdivision (*F*_1,15_ = 9.81, p = 0.007), whereas the main effect of sex (*F*_1,15_ = 0.81, p = 0.383) and the interaction (*F*_1,15_ = 0.26, p = 0.619) were not supported (Fig. 10C). In males, the amplitude of change was higher in the MDmc group than in the MDl group (t = 2.65, p = 0.018), consistent with a stronger aversive sensibility in MDmc. In females, the corresponding MDmc vs MDl comparison did not reach significance (t = 1.80, p = 0.091) despite a similar trend. No sex-specific differences were found in the MDmc (t = 0.97, p = 0.349) and the MDl (t = 0.28, p = 0.780) groups.

Together, these analyses indicate that MDmc and MDl projections to ACC both contribute to the affective-motivational dimension of pain, but in distinct ways. Inhibiting MDmc-ACC selectively increases avoidance of the non-noxious compartment. In contrast, MDl-ACC inhibition does not change PEAP scores despite marked sensory hypersensitivity.

#### 10.2. MD-ACC activation

Next, we evaluated how MD-ACC ChR2 activation shapes the affective-motivational pain component (Fig. 10D). The analysis revealed main effects of optogenetic condition (*F*_(1,32)_ = 6.17, p = 0.020) on the time spent in the lit compartment, sex (*F*_(1,32)_ = 5.06, p = 0.030), and subdivision (*F*_(1,32)_ = 14.65, p < 0.001). The only statistically supported interaction was the light stimulation × subdivision interaction (*F*_(1,32)_ = 19.70, p < 0.001) (Fig. 10E). In males, photostimulation of MDmc-ACC increased the time spent in the light compartment compared with the light off baseline (t = 3.57, p = 0.030). In contrast, MDl-ACC activation did not change light-compartment occupancy (t = 1.19, p = 0.950). In females, the same preferences were observed. MDmc-ACC projections activation increased time in the light compartment compared to the light off baseline (t = 3.36, p = 0.040), while the activation of MDl-ACC projections remained unchanged (t = 0.76, p = 0.980). Normalization of the measures revealed a main effect of subdivision (*F*_(1,16)_ = 11.44, p = 0.004), whereas the effect of sex (*F*_(1,16)_ = 0.00, p = 0.972) and the interaction (*F*_(1,16)_ = 0.23, p = 0.641) were not significant (Fig. 10F). In males, the amplitude of change in light-compartment time was greater in MDmc than MDl animals (t = 2.73, p = 0.015). In females, the same difference was observed but did not reach significance (t = 2.06, p = 0.057).

Taken together, PEAP analyses show that MDmc-ACC activation or inhibition reliably increases time spent in the non-noxious light compartment, indicating that changes in MDmc drive are tightly coupled to affective-motivational escape/avoidance. By contrast, MDl-ACC activation produces no effect.

#### 10.3. MD-PrL inhibition

To assess the affective-motivational component in the PEAP during optogenetic inhibition of MD-PrL projections using ArchT opsin (Fig. 10G), a three-way ANOVA was conducted. This analysis revealed a main effect of optogenetic condition (*F*_(1,32)_ = 84.75, P < 0.001). In contrast, neither sex (*F*_(1,32)_ = 0.38, P = 0.540) nor subdivision (*F*_(1,32)_ = 2.10, P = 0.160) showed a significant main effect. Among the interaction terms, only the optogenetic condition × subdivision interaction reached significance (*F*_(1,32)_ = 5.88, P = 0.020) (Fig. 10H). In males, light delivery increased the time spent in the illuminated compartment in the MDmc condition (t = 6.21, p < 0.001), whereas the change in the MDl condition did not reach significance (t = 2.82, p = 0.100). In females, light delivery increased time spent in the illuminated compartment in both MDmc (t = 5.43, p < 0.001) and MDl conditions (t = 3.96, p = 0.007).

The normalization of time in the lit compartment as a percent change relative to each animal’s light-off level (Fig. 10I) did not reveal an effect of sex (*F*_(1,16)_ = 0.09, p = 0.774), or sex × subdivision interaction (*F*_(1,16)_ = 0.69, p = 0.420). The effect of subdivision showed a trend (*F*_(1,16)_ = 4.46, p = 0.051); however, it did not reach significance. Overall, when expressed relative to light-off values, the modulation of lit-compartment occupancy appeared broadly comparable across sexes and subdivisions.

These results show that both inhibiting MDmc and MDl PrL projections to the PrL increases time spent in the non-noxious light compartment in both sexes.

#### 10.4. MD-PrL activation

The assessment of the affective-motivational component of pain during optogenetic activation of MD-PrL projections (Fig. 10J) revealed a main effect of light stimulation on time in the light compartment (*F*_(1,32)_ = 84.75, p < 0.001). This effect was not driven by sex (*F*_(1,32)_ = 0.38, p = 0.540) or by subdivision alone (*F*_(1,32)_ = 2.10, p = 0.160). Importantly, the impact of light stimulation depended on the MD subdivision (light delivery × subdivision: *F*_(1,32)_ = 5.88, p = 0.020) (Fig. 10K). In males, light delivery increased escape-compartment time in the MDmc-PrL group (t = 6.21, p < 0.001), while in the MDl-PrL group, there was no difference (t = 2.82, p = 0.100). In females, light delivery increased escape-compartment time after MDmc-PrL activation (t = 5.43, p < 0.001) but not after MDl-PrL activation (t = 1.04, p = 0.840).

Normalized responses revealed a main effect of subdivision (*F*_(1,16)_ = 21.62, p < 0.001), with no effect of sex (*F*_(1,16)_ = 0.0001, p = 0.99) or interaction (*F*_(1,16)_ = 0.04, p = 0.831) (Fig. 10L). The effects triggered by MDmc-PrL activation are significantly stronger than in the MDl-PrL group in males (t = 3.44, p = 0.003) and females (t = 3.13, p = 0.006). Sex comparisons further indicated no difference within the MDmc-PrL group (t = 0.14, p = 0.889) or the MDl-PrL group (t = 0.16, p = 0.874).

These PEAP data show that MDmc-PrL activation enhances avoidance of the painful dark compartment. In contrast, MDl-PrL activation induces analgesia in thermal and cold assays and is accompanied by a much weaker or even absent light-ward shift, such that MDl-stimulated animals can remain in the nociceptive dark compartment despite ongoing stimulation. This pattern contrasts with the blunted avoidance seen during MDl-ACC inhibition, and indicates that MDl-PrL projections contribute to the affective-motivational control of pain in a manner that is tightly coupled to their analgesic impact on sensory readouts.

## Discussion

Across lesions, tracing, cortical activity mapping, and bidirectional optogenetic perturbations, our data indicate that MD subdivisions exert dissociable control over pain dimensions by engaging distinct ACC and PrL inhibitory microcircuits. Both MDmc and MDl disruption increased sensory-discriminative nociception, but their effects on affective-motivational behavior diverged, showing that sensory hypersensitivity and pain-related aversion are not strictly coupled. Pathway-specific optogenetic inhibition and activation further showed that MDmc and MDl projections are not simple on/off regulators of pain, but exert target-dependent control over sensory thresholds and aversion. This dissociation was supported anatomically by a subdivision- and target-specific organization of MD-mPFC pathways, with MDl projections preferentially associated with PV-related motifs and MDmc projections more closely linked to SOM-related motifs, particularly in the ACC. Functionally, cFos activation suggests that MD disruption does not produce homogeneous cortical hypoactivity or hyperactivity, but rather rebalances layer-specific excitatory-inhibitory recruitment within ACC and PrL circuits. Together, these findings support a model in which MDmc preferentially couples nociceptive gain to aversive-motivational responding, whereas MDl exerts more flexible, target-dependent control over sensory and affective pain dimensions.

Over the past two decades, converging behavioral and circuit work has established that pain is multidimensional and that prefrontal-cingulate networks are especially important for the affective-motivational component driving avoidance and “suffering-like” behavior [62,27,48]. In rodents, the PEAP was developed to quantify this aversive dimension beyond reflexive thresholds [41,15], and ACC manipulations were shown to blunt escape/avoidance without necessarily normalizing mechanical hypersensitivity, highlighting a partial dissociation between sensory gain and aversive choice [43]. This distinction is further supported by studies showing that chronic pain can drive anxio-depressive-like behaviors through ACC circuits partially independent from peripheral hypersensitivity [79,78], and that ACC synaptic plasticity, particularly long-term potentiation, preferentially contributes to affective pain behaviors without uniformly scaling mechanical thresholds [83,31]. The MD has been framed as a key higher-order hub for mPFC function, with anatomically heterogeneous projections capable of shaping cortical E/I balance by recruiting specific interneuron subpopulations. PV-mediated feedforward inhibition is an example of mechanisms activated by MD projections [63,8,2,24]. Recent work has linked MD-ACC and PrL circuits to pain aversion, hypersensitivity, and GABAergic control of pain-related behavior [81,49,46]. However, most studies have treated the MD as a unitary structure or focused on single pathways, leaving unresolved how MD subdivisions, mPFC targets, and inhibitory microcircuits coordinate or dissociate sensory thresholds from affective-motivational avoidance. In the present study, we used functional tracing to uncover specific MD-mPFC contributions to the association or dissociation of sensory and emotional components of pain.

The dissociation between hypersensitivity and avoidance, particularly after MDl disruption, suggests that MD lesions affect pain through separable sensory and emotional mechanisms. Our data point to a subdivision-specific organization in which MDmc and MDl differentially contribute to the transformation of nociceptive input into aversive motivational value. Consistently, our 3D tracing showed preferential MDl-PV and MDmc-SOM associations in the ACC, with weaker segregation in the PrL, supporting anatomically and functionally differentiated thalamo-cingulate microcircuits [75,47].

SOM interneurons regulate dendritic integration and contextual filtering, whereas PV interneurons provide rapid perisomatic inhibition controlling pyramidal output, synchrony, and temporal precision [71,7,2]. Because SOM interneurons can inhibit PV interneurons, they may also indirectly shape pyramidal output through SOM-PV interactions [58,71]. Within this model, MDmc-ACC projections may support the integration of nociceptive information into an emotional context through SOM-related mechanisms. Their disruption could weaken dendritic filtering and SOM-PV coordination, favoring pyramidal disinhibition, increased cFos, and amplification of both sensory reactivity and aversive value. This is consistent with studies showing that MD-ACC inputs reconfigure ACC microcircuits during pain states and modulate aversion, and that thalamo-cingulate plasticity contributes to hypersensitivity and aversive learning in chronic pain [49,72,46].

By contrast, MDl-ACC disruption may preferentially affect PV-related inhibitory recruitment, which could contribute to altered cortical processing of nociceptive inputs. This would explain why hypersensitivity may occur without a proportional increase in avoidance, consistent with the dissociation between nociceptive reactivity and aversive choice [41,75]. These findings suggest that similar sensory perturbations can acquire different affective-motivational weights depending on which MD-ACC pathway is engaged and which inhibitory circuit is engaged. The increased global cFos supports this hypothesis of reduced PV/SOM recruitment and cortical disinhibition.

At the PrL level, MDmc and MDl effects appeared more convergent. The weaker subdivision-specific segregation of MD projections onto PrL interneuron populations suggests that PrL circuits may not primarily encode aversive valence, but rather translate the ongoing nociceptive state into adaptive avoidance responses [56,52,12].

The sensory-discriminative phenotype can then be interpreted as a convergent consequence of disrupted MD-mPFC regulation. The hypersensitivity observed after MDmc-ACC and MDl-ACC inhibition, as well as after their lesions, suggests that both subdivisions contribute to stabilizing nociceptive thresholds. These convergent effects suggest that MD-ACC inputs provide a common thalamic control over nociceptive gain, whereas the affective-motivational consequences remain subdivision dependent. This interpretation is consistent with the dual role of the ACC in affective evaluation and sensory modulation, as ACC pyramidal activity responds to noxious stimulation, relates to withdrawal thresholds, and can bidirectionally regulate nociceptive responses through local ACC microcircuits involving inhibitory interneurons and excitatory pyramidal neurons [62,27,32,26,16,20,29,49]. A complementary mechanism may operate through the PrL, which filters pain-related information and converts it into adaptive behavioral output rather than encoding nociceptive intensity directly [4,75,52,76,21]. Consistently, PrL GABAergic circuits regulate pyramidal output and nociceptive gain [81], while MD-PrL projections may provide the thalamic drive required to recruit PV/SOM inhibitory control [63,71,21,2]. Thus, reduced MD-PrL drive, e.g., after lesions of MD subdivisions, may weaken inhibitory gating, consistent with increased PrL cFos and reduced PV/SOM recruitment after nociceptive stimulation.

Bidirectional perturbation of MDmc-mPFC pathways may converge on a pro-nociceptive, aversive state by destabilizing MD-prefrontal loop dynamics rather than simply increasing or decreasing activity. A parsimonious explanation for why both silencing and activating MDmc terminals in ACC or PrL produced similar behavioral outcomes is that MDmc operates within recurrent thalamo-cortico-thalamic loops that require appropriately patterned drive. Anatomical and functional evidence support direct excitatory feedback in which MD terminals contact pyramidal cells, including deep-layer corticothalamic neurons, enabling reverberatory MD-mPFC interactions [37,40,60,1,68,69]. Thus, perturbing MDmc input may disrupt the balance between thalamic excitation and cortical inhibition, altering the effective output of ACC and PrL circuits involved in pain regulation [14,49]. In this framework, photoinhibition/stimulation may disrupt a stabilizing MDmc drive normally required for MD-prefrontal coordination and pain-modulatory control. This U-shaped model is consistent with evidence that opposite perturbations can produce convergent behavioral effects when they disturb physiological activity patterns rather than scale circuit activity linearly [59,45,49].

A notable strength of this study is that it considers possible sex differences in MD-mPFC pain control. Although sex-related differences in nociceptive sensitivity are well documented [53,67], our data do not support a fundamentally distinct organization of MD-dependent pain regulation in males and females. MD subdivision lesions produced broadly comparable behavioral outcomes across sexes, suggesting that MD-mPFC contributions to sensory gain and affective-motivational aversion are largely conserved across sexes. Some sex-linked variations were observed in PV/SOM recruitment and laminar cFos profiles, indicating subtle differences in inhibitory microcircuit engagement, but these did not translate into qualitatively different pain phenotypes.

Importantly, human neuroimaging evidence suggests that the MD is part of altered brain networks in chronic pain [34,30,25]. These networks include medial prefrontal regions, and stronger MD-prefrontal functional connectivity has been associated with higher negative affect in patients [25]. In this context, our results suggest that MD-PFC control is not uniform and indicate a higher complexity depending on specific pathways connecting MD subdivisions and cortical targets. This circuit specificity also supports a cautious therapeutic perspective: rather than broadly activating or silencing prefrontal regions, more targeted strategies may aim to restore appropriate mPFC circuit function within defined pathways. This view is consistent with current calls to target prefrontal dysfunction in chronic pain, while emphasizing that therapeutic effects may depend on whether the circuit is hypoactive or hyperactive and whether the intervention is applied in a normal or pain-altered state [49,35].

## Limitations and future directions

Although our approach discriminated between MD projections, it did not isolate finer subdivision microdomains within MD (e.g., separating MDm from MDc). Future work using higher-resolution anatomical definitions could refine whether MDmc-related phenotypes reflect partially distinct medial and central subcircuits. In addition, our optogenetic manipulations provided robust pathway-level control but were not designed to isolate specific neuronal populations within the MD or the target cortical regions. Consequently, although our anatomical and activity-based analyses point to laminar and PV/SOM-related microcircuit engagement, they do not establish the causal contribution of a single interneuron class or cortical layer. Cell-type- and layer-restricted manipulations will therefore be necessary to determine how distinct MD-prefrontal microcircuits contribute to sensory-discriminative and affective-motivational dimensions of pain.

## CRediT authorship contribution statement

Hanane Iben-Daoudi: Conceptualization, Methodology, Formal analysis, Investigation, Data curation, Writing - original draft, Visualization. Saadia Ba-M’hamed: Conceptualization, Methodology, Validation, Formal analysis, Writing - review and editing. Fatima-Zahra Lamghari Moubarrad: Conceptualization, Methodology, Validation, Resources, Writing - review and editing, Supervision. Mohamed Bennis: Conceptualization, Methodology, Validation, Resources, Writing - review and editing, Visualization, Supervision. Marc Landry: Conceptualization, Methodology, Supervision, Writing - review and editing, Visualization, Resources, Validation, Funding acquisition. Zakaria Ouhaz: Conceptualization, Methodology, Writing - review and editing, Visualization, Supervision.

## Acknowledgments

We would like to thank Franck Aby for his support with the optogenetic experiments. We are also grateful to Thibault Dhellemmes for his assistance with BioLoc3D analyses during the tracing experiments and to Sandra Dovero for her help with the slide-scanner acquisitions for the immunohistochemical experiments. We also thank the Broca Center animal experimentation facility (PIV).

## Data Availability Statement

The data supporting the findings of this study are available in an online repository. The repository can be found below: https://github.com/marclandry33/MD-PFC-pathways-in-pain

## Abbreviation list

AAV: Adeno-associated virus
ACC: Anterior cingulate cortex
ANOVA: Analysis of variance
AP: Anteroposterior coordinate
APAFIS: French ethical authorization/application system number
ArchT: Archaerhodopsin-T
BSA: Bovine serum albumin
ChR2: Channelrhodopsin-2
CM: Central medial nucleus
CL: Central lateral nucleus
ContraL: Contralateral hind paw
D3V: Dorsal third ventricle
DAPI: 4′,6-diamidino-2-phenylindole
DV: Dorsoventral coordinate
EGFP: Enhanced green fluorescent protein
EYFP: Enhanced yellow fluorescent protein
H134R: ChR2 point mutation used in the hChR2(H134R) construct
Hz: Hertz
ID: Inner diameter
IL: Infralimbic cortex
IMD: Intermediodorsal nucleus
i.p.: Intraperitoneal
IpsiL: Ipsilateral hind paw
L1: Cortical layer 1
L2/3: Cortical layers 2/3
L5: Cortical layer 5
L6: Cortical layer 6
LC: Lucent connector
LHb: Lateral habenula
M2: Secondary motor cortex
MD: Mediodorsal thalamus
MDc: Central mediodorsal thalamic subdivision
MDl: Lateral mediodorsal thalamic subdivision
MDm: Medial mediodorsal thalamic subdivision
MDmc: Medial-central mediodorsal thalamic subdivision
MGV: Mean gray value
MHb: Medial habenula
ML: Mediolateral coordinate
mPFC: Medial prefrontal cortex
NA: Numerical aperture
NMDA: N-methyl-D-aspartate
PBS: Phosphate-buffered saline
PC: Paracentral nucleus
PEAP: Place escape/avoidance paradigm
PFA: Paraformaldehyde
PFC: Prefrontal cortex
PrL: Prelimbic cortex
PV: Parvalbumin
PVP: Paraventricular posterior nucleus
ROI / ROIs: Region of interest/regions of interest
s.c.: Subcutaneous
SEM: Standard error of the mean
SOM: Somatostatin
vg/mL: Viral genomes per milliliter

---

**Figure S1. Histological verification and quantification of subdivision-selective excitotoxic lesions in the mediodorsal thalamus (MDmc vs MDl).** (A) Schematic reconstructions of the maximal lesion spread (colored overlays) across rostro-caudal anteroposterior (AP) coordinates (mm, relative to bregma) for the MDmc (top row) and MDl (bottom row) lesion groups. Colored regions indicate the estimated area of neuronal loss within the targeted MD subdivision across animals. (B) High-magnification Nissl-stained micrographs illustrating cellular architecture within MDm, MDc, and MDl for Sham, MDmc lesion, and MDl lesion animals. Black arrowheads indicate examples of Nissl-stained neuronal profiles used for density quantification (and highlight the reduction of identifiable neuronal somata in the targeted subdivision). Scale bar: 100µm. (C) Low-magnification thalamic sections showing the global lesion site and extent at selected AP levels for Sham, MDmc lesion, and MDl lesion groups. Scale bar: 300µm. (D) Quantification of neuronal density and lesion selectivity within MD subdivisions; Left: Neuronal cell density (neurons/mm²) in MDm, MDc, and MDl across groups; Right: Corresponding percent cell loss (normalized to Sham) for each lesion group, highlighting preferential loss in the targeted subdivision while sparing non-targeted MD regions. (E) Percent cell loss (relative to Sham) in surrounding thalamic nuclei to evaluate lesion specificity. Data are presented as mean ± SEM (n = 8). Data analyzed using one-way ANOVA followed by Holm-Šidák multiple-comparison *post-hoc* tests or t-tests. (***p < 0.001 Lesioned subdivision vs. Sham subdivision; ###p < 0.001 MDmc vs MDl). CL, centrolateral; CM, centromedial; PV, paraventricular; LHeb, lateral habenula; PC, paracentral.

**Figure S2. GFP control experiments show that optical stimulation of MDmc/MDl-ACC pathways does not alter mechanical, thermal, or affective pain behaviors in male or female rats.** (A) Schematic illustration of the control experiment in which AAV5-CaMKIIα-GFP (without opsin) was injected into the MDmc and an optic fibre was implanted over the ACC. (B) paw-withdrawal thresholds (g) of the ipsilateral (IpsiL) and contralateral (ContraL) hind paws in the Von Frey test. (C,D) Hot-plate (C) and cold-plate (D) tests for the same MDmc-ACC GFP cohort, showing paw-withdrawal latencies (s). (E) Schematic of the analogous control experiment with AAV5-CaMKIIα-GFP injection into the MDl and an optic fibre over the ACC. (F) paw-withdrawal thresholds (g) in the Von Frey test. (G, H) Hot-plate (G) and cold-plate (H) paw-withdrawal latencies. (I) Schematic of the place escape/avoidance paradigm (PEAP) control experiment in which GFP-only virus was injected into MDmc or MDl, and an optic fibre was placed over the ACC. (J) Time spent on the light side (%) during PEAP. Data are presented as mean ± SEM. Sample sizes were as follows: MDmc-ACC-GFP (males n = 6, females n = 5) and MDl-PrL-GFP (males n = 5, females n = 6). For Von Frey, hot-plate, and cold-plate tests, repeated-measures two-way ANOVA was performed with stage and sex as factors. For PEAP, three-way ANOVA with factors subdivision, light condition, and sex was used.

**Figure S3. GFP control experiments show that optical stimulation of MDmc/MDl-PrL pathways does not alter mechanical, thermal, or affective pain behaviors in male or female rats.** (A) Schematic illustration of the control experiment in which AAV5-CaMKIIα-GFP (without opsin) was injected into the MDmc and an optic fibre was implanted over the PrL. (B) paw-withdrawal thresholds (g) of the ipsilateral (IpsiL) and contralateral (ContraL) hind paws in the Von Frey test. (C, D) Hot-plate (C) and cold-plate (D) tests for the same MDmc-PrL GFP cohort, showing paw-withdrawal latencies (s). (E) Schematic of the analogous control experiment with AAV5-CaMKIIα-GFP injection into the MDl and an optic fibre over the PrL. (F) paw-withdrawal thresholds (g) in the Von Frey test. (G, H) Hot-plate (G) and cold-plate (H) paw-withdrawal latencies. (I) Schematic of PEAP control experiment in which GFP-only virus was injected into MDmc or MDl, and an optic fibre was placed over the PrL. (J) Time spent on the light side (%) during PEAP. Data are presented as mean ± SEM. Sample sizes were as follows: MDmc-PrL-GFP (males n = 5, females n = 5) and MDl-PrL-GFP (males n = 6, females n = 6).. For Von Frey, hot-plate, and cold-plate tests, repeated-measures two-way ANOVA was performed with stage and sex as factors. For PEAP, three-way ANOVA with factors subdivision, light condition, and sex was used.

**Figure S4. Spatial mapping of the optogenetically driven cFos response around the fiber-optic tip in ACC and PrL cortex following MD subdivision-specific terminal modulation.** cFos positive cell density (cells/mm²) was quantified as a function of distance from the fiber-optic tip (0-1200 µm; pooled in 300-µm distance bins). Across conditions, cFos modulation was maximal in proximity to the fiber tip and progressively decayed with distance, indicating a spatially restricted optogenetic footprint. The ACC panels depict the distance-dependent cFos profile produced by MDmc-ACC and MDl-ACC terminal photoinhibition (ArchT) (A-B) and photoactivation (ChR2) (C-D) (separate plots for each MD subdivision and opsin). The PrL panels show the corresponding distance-dependent cFos profiles for MDmc-PrL and MDl-PrL terminal photoinhibition (ArchT) (E-F) and photoactivation (ChR2) (G-H). Data are presented as mean ± SEM (n = 4 per group) and were analyzed using a one-way ANOVA followed by Holm-Šidák *post-hoc* tests (*p < 0.05; **p < 0.01; ***p < 0.001 relative to the data point at 1200 µm).

## References

[1] Agmon A, Connors BW. Thalamocortical responses of mouse somatosensory (barrel) cortex in vitro. Neuroscience 1991;41:365–379. doi:10.1016/0306-4522(91)90333-J

[2] Anastasiades PG, Carter AG. Circuit organization of the rodent medial prefrontal cortex. Trends Neurosci 2021;44:550–563. doi:10.1016/j.tins.2021.03.006

[3] Berendse HW, Groenewegen HJ. Restricted cortical termination fields of the midline and intralaminar thalamic nuclei in the rat. Neuroscience 1991;42:73–102. doi:10.1016/0306-4522(91)90151-D

[4] Bush G, Luu P, Posner MI. Cognitive and emotional influences in anterior cingulate cortex. Trends Cogn Sci 2000;4:215–222. doi:10.1016/S1364-6613(00)01483-2

[5] Bushnell MC, Čeko M, Low LA. Cognitive and emotional control of pain and its disruption in chronic pain. Nat Rev Neurosci 2013;14:502–511. doi:10.1038/nrn3516

[6] Chen T, Taniguchi W, Chen QY, Tozaki-Saitoh H, Song Q, Liu RH, Koga K, Matsuda T, Kaito-Sugimura Y, Wang J, Li ZH, Lu YC, Inoue K, Tsuda M, Li YQ, Nakatsuka T, Zhuo M. Top-down descending facilitation of spinal sensory excitatory transmission from the anterior cingulate cortex. Nat Commun 2018;9:1886. doi:10.1038/s41467-018-04309-2

[7] Cichon J, Blanck TJJ, Gan WB, Yang G. Activation of cortical somatostatin interneurons prevents the development of neuropathic pain. Nat Neurosci 2017;20:1122–1132. doi:10.1038/nn.4595

[8] Delevich K, Tucciarone J, Huang ZJ, Li B. The mediodorsal thalamus drives feedforward inhibition in the anterior cingulate cortex via parvalbumin interneurons. J Neurosci 2015;35:5743–5753. doi:10.1523/JNEUROSCI.4565-14.2015

[9] Deuis JR, Dvorakova LS, Vetter I. Methods used to evaluate pain behaviors in rodents. Front Mol Neurosci 2017;10:284. doi:10.3389/fnmol.2017.00284

[10] Dhellemmes T, Inet R, Cordelières F, Landry M. BioLoc3D (B3D): Protocol for 3D automated analysis of colocalization in microscopy. protocols.io 2025;version 2. doi:10.17504/protocols.io.kxygx4p5zl8j/v2

[11] Dong WK, Ryu H, Wagman IH. Nociceptive responses of neurons in medial thalamus and their relationship to spinothalamic pathways. J Neurophysiol 1978;41:1592–1613. doi:10.1152/jn.1978.41.6.1592

[12] Drake RA, Steel KA, Apps R, Lumb BM, Pickering AE. Loss of cortical control over the descending pain modulatory system determines the development of the neuropathic pain state in rats. eLife 2021;10:e65156. doi:10.7554/eLife.65156

[13] Espejo EF, Mir D. Structure of the rat’s behavior in the hot plate test. Behav Brain Res 1993;56:171–176. doi:10.1016/0166-4328(93)90035-O

[14] Floresco SB, Grace AA. Gating of hippocampal-evoked activity in prefrontal cortical neurons by inputs from the mediodorsal thalamus and ventral tegmental area. J Neurosci 2003;23:3930–3943. doi:10.1523/JNEUROSCI.23-09-03930.2003

[15] Fuchs PN, McNabb CT. The place escape/avoidance paradigm: A novel method to assess nociceptive processing. J Integr Neurosci 2012;11:61–72. doi:10.1142/S0219635212500045

[16] Fuchs PN, Peng YB, Boyette-Davis JA, Uhelski ML. The anterior cingulate cortex and pain processing. Front Integr Neurosci 2014;8:35. doi:10.3389/fnint.2014.00035

[17] García-Cabezas MÁ, John YJ, Barbas H, Zikopoulos B. Distinction of neurons, glia and endothelial cells in the cerebral cortex: An algorithm based on cytological features. Front Neuroanat 2016;10:107. doi:10.3389/fnana.2016.00107

[18] Giesler GJ Jr, Menétrey D, Basbaum AI. Differential origins of spinothalamic tract projections to medial and lateral thalamus in the rat. J Comp Neurol 1979;184:107–126. doi:10.1002/cne.901840107

[19] Groenewegen HJ. Organization of the afferent connections of the mediodorsal thalamic nucleus in the rat, related to the mediodorsal-prefrontal topography. Neuroscience 1988;24:379–431. doi:10.1016/0306-4522(88)90339-9

[20] Gu L, Uhelski ML, Anand S, Romero-Ortega M, Kim YT, Fuchs PN, Mohanty SK. Pain inhibition by optogenetic activation of specific anterior cingulate cortical neurons. PLoS One 2015;10:e0117746. doi:10.1371/journal.pone.0117746

[21] Halassa MM, Kastner S. Thalamic functions in distributed cognitive control. Nat Neurosci 2017;20:1669–1679. doi:10.1038/s41593-017-0020-1

[22] Hsu MM, Kung JC, Shyu BC. Evoked responses of the anterior cingulate cortex to stimulation of the medial thalamus. Chin J Physiol 2000;43:81–89.

[23] Hsu MM, Shyu BC. Electrophysiological study of the connection between medial thalamus and anterior cingulate cortex in the rat. Neuroreport 1997;8:2701–2707. doi:10.1097/00001756-199708180-00013

[24] Iben-Daoudi H, Bennis M, Landry M, Lamghari FZ, Ba-M’hamed S, Ouhaz Z. Differential Implications of the mediodorsal thalamic nucleus subdivisions in regulating prefrontal cortex GAD67 and GABAB receptors expression: behavioral and cognitive outcomes. Neuroscience 2026;594:203–223. doi:10.1016/j.neuroscience.2025.12.024

[25] Iwabuchi SJ, Drabek MM, Cottam WJ, Tadjibaev A, Mohammadi-Nejad AR, Sotiropoulos S, Fernandes GS, Valdes AM, Zhang W, Doherty M, Walsh DA, Auer DP. Medio-dorsal thalamic dysconnectivity in chronic knee pain: A possible mechanism for negative affect and pain comorbidity. Eur J Neurosci 2023;57:373–387. doi:10.1111/ejn.15880

[26] Iwata K, Kamo H, Ogawa A, Tsuboi Y, Noma N, Mitsuhashi Y, Taira M, Koshikawa N, Kitagawa J. Anterior cingulate cortical neuronal activity during perception of noxious thermal stimuli in monkeys. J Neurophysiol 2005;94:1980–1991. doi:10.1152/jn.00190.2005

[27] Johansen JP, Fields HL, Manning BH. The affective component of pain in rodents: Direct evidence for a contribution of the anterior cingulate cortex. Proc Natl Acad Sci U S A 2001;98:8077–8082. doi:10.1073/pnas.141218998

[28] Journée SH, Mathis VP, Fillinger C, Veinante P, Yalcin I. Janus effect of the anterior cingulate cortex: Pain and emotion. Neurosci Biobehav Rev 2023;153:105362. doi:10.1016/j.neubiorev.2023.105362

[29] Kang SJ, Kwak C, Lee J, Sim SE, Shim J, Choi T, Collingridge GL, Zhou M, Kaang BK. Bidirectional modulation of hyperalgesia via the specific control of excitatory and inhibitory neuronal activity in the ACC. Mol Brain 2015;8:81. doi:10.1186/s13041-015-0170-6

[30] Karunakaran KD, Yuan R, He J, Zhao J, Cui JL, Zang YF, Zhang Z, Alvarez TL, Biswal BB. Resting-state functional connectivity of the thalamus in complete spinal cord injury. Neurorehabil Neural Repair 2020;34:122–133. doi:10.1177/1545968319893299

[31] Koga K, Descalzi G, Chen T, Ko HG, Lu J, Li S, Son J, Kim TH, Kwak C, Huganir RL, Zhao MG, Kaang BK, Collingridge GL, Zhuo M. Coexistence of two forms of LTP in ACC provides a synaptic mechanism for the interactions between anxiety and chronic pain. Neuron 2015;85:377–389. doi:10.1016/j.neuron.2014.12.021

[32] Koyama T, Kato K, Tanaka YZ, Mikami A. Anterior cingulate activity during pain-avoidance and reward tasks in monkeys. Neurosci Res 2001;39:421–430. doi:10.1016/S0168-0102(01)00197-3

[33] Krettek JE, Price JL. The cortical projections of the mediodorsal nucleus and adjacent thalamic nuclei in the rat. J Comp Neurol 1977;171:157–192. doi:10.1002/cne.901710204

[34] Kucyi A, Moayedi M, Weissman-Fogel I, Goldberg MB, Freeman BV, Tenenbaum HC, Davis KD. Enhanced medial prefrontal-default mode network functional connectivity in chronic pain and its association with pain rumination. J Neurosci 2014;34:3969–3975. doi:10.1523/JNEUROSCI.5055-13.2014

[35] Kummer K, Sheets PL. Targeting prefrontal cortex dysfunction in pain. J Pharmacol Exp Ther 2024;389:268–276. doi:10.1124/jpet.123.002046

[36] Kuramoto E, Pan S, Furuta T, Tanaka YR, Iwai H, Yamanaka A, Ohno S, Kaneko T, Goto T, Hioki H. Individual mediodorsal thalamic neurons project to multiple areas of the rat prefrontal cortex: A single neuron-tracing study using virus vectors. J Comp Neurol 2017;525:166–185. doi:10.1002/cne.24054

[37] Kuroda M, Murakami K, Oda S, Shinkai M, Kishi K. Direct synaptic connections between thalamocortical axon terminals from the mediodorsal thalamic nucleus (MD) and corticothalamic neurons to MD in the prefrontal cortex. Brain Res 1993;612:339–344. doi:10.1016/0006-8993(93)91683-J

[38] Kuroda M, Murakami K, Shinkai M, Ojima H, Kishi K. Electron microscopic evidence that axon terminals from the mediodorsal thalamic nucleus make direct synaptic contacts with callosal cells in the prelimbic cortex of the rat. Brain Res 1995;677:348–353. doi:10.1016/0006-8993(95)00192-S

[39] Kuroda M, Ojima H, Igarashi H, Murakami K, Okada A, Shinkai M. Synaptic relationships between axon terminals from the mediodorsal thalamic nucleus and layer III pyramidal cells in the prelimbic cortex of the rat. Brain Res 1996;708:185–190. doi:10.1016/0006-8993(95)01438-1

[40] Kuroda M, Yokofujita J, Murakami K. An ultrastructural study of the neural circuit between the prefrontal cortex and the mediodorsal nucleus of the thalamus. Prog Neurobiol 1998;54:417–458. doi:10.1016/S0301-0082(97)00070-1

[41] LaBuda CJ, Fuchs PN. A behavioral test paradigm to measure the aversive quality of inflammatory and neuropathic pain in rats. Exp Neurol 2000;163:490–494. doi:10.1006/exnr.2000.7395

[42] LaGraize SC, Fuchs PN. GABAA but not GABAB receptors in the rostral anterior cingulate cortex selectively modulate pain-induced escape/avoidance behavior. Exp Neurol 2007;204:182–194. doi:10.1016/j.expneurol.2006.10.007

[43] LaGraize SC, LaBuda CJ, Rutledge MA, Jackson RL, Fuchs PN. Differential effect of anterior cingulate cortex lesion on mechanical hypersensitivity and escape/avoidance behavior in an animal model of neuropathic pain. Exp Neurol 2004;188:139–148. doi:10.1016/j.expneurol.2004.04.003

[44] Lançon K, Tian J, Bach H, Drapeau P, Poulin JF, Séguéla P. Synergistic deficits in parvalbumin interneurons and dopamine signaling drive ACC dysfunction in chronic pain. Proc Natl Acad Sci U S A 2025;122:e2502558122. doi:10.1073/pnas.2502558122

[45] Li N, Chen S, Guo ZV, Chen H, Huo Y, Inagaki HK, Chen G, Davis C, Hansel D, Guo C, Svoboda K. Spatiotemporal constraints on optogenetic inactivation in cortical circuits. eLife 2019;8:e48622. doi:10.7554/eLife.48622

[46] Lian YN, Cao XW, Wu C, Pei CY, Liu L, Zhang C, Li XY. Deconstruction the feedforward inhibition changes in the layer III of anterior cingulate cortex after peripheral nerve injury. Commun Biol 2024;7:1237. doi:10.1038/s42003-024-06849-4

[47] Likhtik E, Stujenske JM, Topiwala MA, Harris AZ, Gordon JA. Prefrontal entrainment of amygdala activity signals safety in learned fear and innate anxiety. Nat Neurosci 2014;17:106–113. doi:10.1038/nn.3582

[48] Liu CH, Li HY, Li KS, Li YT, Cheng JB, Wang XH, Huang DX, Yang XF, Ma RQ, Lu XM, Zhu XY. Involvement of mediodorsal thalamus and its related neural circuit in pain regulation in mice. Neurobiol Dis 2025;200:107056. doi:10.1016/j.nbd.2025.107056

[49] Meda KS, Patel T, Braz JM, Malik R, Turner ML, Seifikar H, Basbaum AI, Sohal VS. Microcircuit mechanisms through which mediodorsal thalamic input to anterior cingulate cortex exacerbates pain-related aversion. Neuron 2019;102:944–959.e3. doi:10.1016/j.neuron.2019.03.042

[50] Melzack R, Casey KL. Sensory, motivational, and central control determinants of pain: A new conceptual model. In: Kenshalo DR, editor. The Skin Senses. Springfield, IL: Charles C Thomas, 1968. pp. 423-439.

[51] Miller EK, Cohen JD. An integrative theory of prefrontal cortex function. Annu Rev Neurosci 2001;24:167–202. doi:10.1146/annurev.neuro.24.1.167

[52] Mitchell AS. The mediodorsal thalamus as a higher order thalamic relay nucleus important for learning and decision-making. Neurosci Biobehav Rev 2015;54:76–88. doi:10.1016/j.neubiorev.2015.03.001

[53] Mogil JS. Sex differences in pain and pain inhibition: Multiple explanations of a controversial phenomenon. Nat Rev Neurosci 2012;13:859–866. doi:10.1038/nrn3360

[54] Naliboff BD, Berman S, Chang L, Derbyshire SW, Suyenobu B, Vogt BA, Mandelkern M, Mayer EA. Sex-related differences in IBS patients: central processing of visceral stimuli. Gastroenterology 2003;124:1738–1747. doi:10.1016/S0016-5085(03)00400-1

[55] Nicotra L, Tuke J, Grace PM, Rolan PE, Hutchinson MR. Sex differences in mechanical allodynia: How can it be preclinically quantified and analyzed? Front Behav Neurosci 2014;8:40. doi:10.3389/fnbeh.2014.00040

[56] Parnaudeau S, O’Neill PK, Bolkan SS, Ward RD, Abbas AI, Roth BL, Balsam PD, Gordon JA, Kellendonk C. Inhibition of mediodorsal thalamus disrupts thalamofrontal connectivity and cognition. Neuron 2013;77:1151–1162. doi:10.1016/j.neuron.2013.01.038

[57] Paxinos G, Watson C. The Rat Brain in Stereotaxic Coordinates, sixth ed. San Diego: Academic Press, 2008.

[58] Pfeffer CK, Xue M, He M, Huang ZJ, Scanziani M. Inhibition of inhibition in visual cortex: The logic of connections between molecularly distinct interneurons. Nat Neurosci 2013;16:1068–1076. doi:10.1038/nn.3446

[59] Phillips EAK, Hasenstaub AR. Asymmetric effects of activating and inactivating cortical interneurons. eLife 2016;5:e18383. doi:10.7554/eLife.18383

[60] Pirot S, Jay TM, Glowinski J, Thierry AM. Anatomical and electrophysiological evidence for an excitatory amino acid pathway from the thalamic mediodorsal nucleus to the prefrontal cortex in the rat. Eur J Neurosci 1994;6:1225–1234. doi:10.1111/j.1460-9568.1994.tb00621.x

[61] Price DD. Psychological and neural mechanisms of the affective dimension of pain. Science 2000;288:1769–1772. doi:10.1126/science.288.5472.1769

[62] Rainville P, Duncan GH, Price DD, Carrier B, Bushnell MC. Pain affect encoded in human anterior cingulate but not somatosensory cortex. Science 1997;277:968–971. doi:10.1126/science.277.5328.968

[63] Rotaru DC, Barrionuevo G, Sesack SR. Mediodorsal thalamic afferents to layer III of the rat prefrontal cortex: Synaptic relationships to subclasses of interneurons. J Comp Neurol 2005;490:220–238. doi:10.1002/cne.20661

[64] Sellmeijer J, Mathis V, Hugel S, Li XH, Song Q, Chen QY, Barthas F, Lutz PE, Karatas M, Luthi A, Veinante P, Barrot M, Zhuo M, Yalcin I. Hyperactivity of anterior cingulate cortex areas 24a/24b drives chronic pain-induced anxiodepressive-like consequences. J Neurosci 2018;38:3102–3115. doi:10.1523/JNEUROSCI.3195-17.2018

[65] Shackman AJ, Salomons TV, Slagter HA, Fox AS, Winter JJ, Davidson RJ. The integration of negative affect, pain and cognitive control in the cingulate cortex. Nat Rev Neurosci 2011;12:154–167. doi:10.1038/nrn2994

[66] Sherman SM, Guillery RW. The role of the thalamus in the flow of information to the cortex. Philos Trans R Soc Lond B Biol Sci 2002;357:1695–1708. doi:10.1098/rstb.2002.1161

[67] Sorge RE, Totsch SK. Sex differences in pain. J Neurosci Res 2017;95:1271–1281. doi:10.1002/jnr.23841

[68] Swadlow HA. Thalamocortical control of feed-forward inhibition in awake somatosensory “barrel” cortex. Philos Trans R Soc Lond B Biol Sci 2002;357:1717–1727. doi:10.1098/rstb.2002.1156

[69] Swadlow HA. Fast-spike interneurons and feedforward inhibition in awake sensory neocortex. Cereb Cortex 2003;13:25–32. doi:10.1093/cercor/13.1.25

[70] Tracey I, Mantyh PW. The cerebral signature for pain perception and its modulation. Neuron 2007;55:377–391. doi:10.1016/j.neuron.2007.07.012

[71] Tremblay R, Lee S, Rudy B. GABAergic interneurons in the neocortex: From cellular properties to circuits. Neuron 2016;91:260–292. doi:10.1016/j.neuron.2016.06.033

[72] Valentinova K, Acuña MA, Ntamati NR, Nevian NE, Nevian T. An amygdala-to-cingulate cortex circuit for conflicting choices in chronic pain. Cell Rep 2023;42:113125. doi:10.1016/j.celrep.2023.113125

[73] Van der Werf YD, Witter MP, Groenewegen HJ. The intralaminar and midline nuclei of the thalamus. Anatomical and functional evidence for participation in processes of arousal and awareness. Brain Res Rev 2002;39:107–140. doi:10.1016/S0165-0173(02)00181-9

[74] Vertes RP. Differential projections of the infralimbic and prelimbic cortex in the rat. Synapse 2004;51:32–58. doi:10.1002/syn.10279

[75] Vogt BA. Pain and emotion interactions in subregions of the cingulate gyrus. Nat Rev Neurosci 2005;6:533–544. doi:10.1038/nrn1704

[76] Wang GQ, Cen C, Li C, Cao S, Wang N, Zhou Z, Liu XM, Xu Y, Tian NX, Zhang Y, Wang J, Wang LP, Wang Y. Deactivation of excitatory neurons in the prelimbic cortex via Cdk5 promotes pain sensation and anxiety. Nat Commun 2015;6:7660. doi:10.1038/ncomms8660

[77] Wolff M, Vezoli J, Procyk E. Executive control by the mediodorsal thalamus. Neuron 2024;112:509–525. doi:10.1016/j.neuron.2024.01.002

[78] Yalcin I, Barrot M. The anxiodepressive comorbidity in chronic pain. Curr Opin Anaesthesiol 2014;27:520–527. doi:10.1097/ACO.0000000000000116

[79] Yalcin I, Bohren Y, Waltisperger E, Sage-Ciocca D, Yin JC, Freund-Mercier MJ, Barrot M. A time-dependent history of mood disorders in a murine model of neuropathic pain. Biol Psychiatry 2011;70:946–953. doi:10.1016/j.biopsych.2011.07.017

[80] Yamamura H, Iwata K, Tsuboi Y, Toda K, Kitajima K, Shimizu N, Nomura H, Hibiya J, Fujita S, Sumino R. Morphological and electrophysiological properties of ACCx nociceptive neurons in rats. Brain Res 1996;735:83–92. doi:10.1016/0006-8993(96)00561-6

[81] Zhang Z, Gadotti VM, Chen L, Souza IA, Stemkowski PL, Zamponi GW. Role of prelimbic GABAergic circuits in sensory and emotional aspects of neuropathic pain. Cell Rep 2015;12:752–759. doi:10.1016/j.celrep.2015.07.001

[82] Zhang Z, Zhang Y, Si J, Mao H, He F, Ning J, Ma G, Chen X, Xiao H, Zhu Y, Zhang H, Lu Y, Liu Q, Nian M, Sun S, Yu S, Fan Z, Jin Z, Huang J. Synaptic dysfunction in the anterior cingulate cortex underlies pain-anxiety comorbidity in a mandibular asymmetry mouse model. Adv Sci 2025;12:e09509. doi:10.1002/advs.202509509

[83] Zhuo M. Cortical excitation and chronic pain. Trends Neurosci 2008;31:199-207. doi:10.1016/j.tins.2008.01.003

[84] Zubieta JK, Smith YR, Bueller JA, Xu Y, Kilbourn MR, Jewett DM, Meyer CR, Koeppe RA, Stohler CS. Mu-opioid receptor-mediated antinociceptive responses differ in men and women. J Neurosci 2002;22:5100–5107. doi:10.1523/JNEUROSCI.22-12-05100.2002

