## Supplementary material for "Rat mediodorsal thalamic subdivisions differentially modulate the sensory and affective components of pain through distinct prefrontal pathways"

| | | | cFos Density (cFos+/mm$^2$) (**Mean ± SEM**) LC groups | | | Two-way ANOVA | | | Pairwise comparisons within MD LC | | | Pairwise comparisons within sex | | |
|---|---|---|---|---|---|---|---|---|---|---|---|---|---|---|
| PFC sub-regions | Layers | Sex | MD sham | MDmc | MDl | LC $F_{(2,37)}$ | Sex $F_{(1,37)}$ | LC x Sex $F_{(2,37)}$ | t1 | t2 | t3 | t4 | t5 | t6 |
| ACC | 1 | M | 11.97 ± 6.19 | 9.52 ± 4.93 | 10.16 ± 6.11 | 0.45 | 0.02 | 0.14 | 0.29 | 0.20 | 0.07 | 0.51 | 0.09 | 0.18 |
| | | F | 16.32 ± 8.04 | 8.70 ± 5.10 | 8.57 ± 5.07 | | | | 0.94 | 0.89 | 0.01 | | | |
| | 2/3 | M | 519.42 ± 31.31 | 744.32 ± 27.28 | 966.55 ± 26.26 | 66.98*** | 0.21 | 1.49 | 4.40*** | 8.15*** | 4.17*** | 0.04 | 1.64 | 0.74 |
| | | F | 516.94 ± 20.69 | 828.03 ± 57.25 | 926.65 ± 41.98 | | | | 5.90*** | 8.03*** | 1.93 | | | |
| | 5 | M | 629.94 ± 28.59 | 467.18 ± 27.66 | 783.61 ± 39.78 | 40.35*** | 14.31*** | 0.80 | 3.78** | 3.32** | 7.05*** | 1.88 | 1.49 | 3.13** |
| | | F | 546.12 ± 28.49 | 402.45 ± 26.10 | 642.74 ± 34.93 | | | | 3.23** | 2.24* | 5.58*** | | | |
| | 6 | M | 410,18 ± 48,60 | 400.73 ± 34.89 | 456.10 ± 32.43 | 0.58 | 0.19 | 0.44 | 0.15 | 0.68 | 0.58 | 0.99 | 0.07 | 0.30 |
| | | F | 473.75 ± 59.99 | 405.13 ± 45.82 | 435.98 ± 39.91 | | | | 1.07 | 0.60 | 0.50 | | | |
| PrL | 1 | M | 14.14 ± 4.52 | 11 .42 ± 3.52 | 13.97 ± 4.58 | 0.08 | 1.35 | 0.02 | 0.38 | 0.02 | 0.34 | 0.60 | 0.85 | 0.56 |
| | | F | 18.50 ± 6.19 | 17.42 ± 4.91 | 18.09 ± 5.92 | | | | 0.15 | 0.05 | 0.09 | | | |
| | 2/3 | M | 527.04 ± 43.94 | 759.62 ± 58.63 | 684.58 ± 35.65 | 2.85 | 4.25* | 4.67 | 3.79** | 2.38* | 1.17 | 1.002 | 3.35** | 1.28 |
| | | F | 590.60 ± 17.71 | 553.84 ± 45.81 | 602.31 ± 43.61 | | | | 0.58 | 0.19 | 0.79 | | | |
| | 5 | M | 576.60 ± 19.56 | 394.66 ± 36.81 | 576.60 ± 19.56 | 14.48*** | 0.27 | 6.38** | 1.74 | 1.83 | 3.57** | 2.51* | 1.06 | 2.35* |
| | | F | 607.14 ± 28.16 | 342.90 ± 32.03 | 456.40 ± 37.36 | | | | 5.23*** | 3.08** | 2.32* | | | |
| | 6 | M | 468.68 ± 46.97 | 394.28 ± 38.57 | 538.43 ± 30.21 | 2.66 | 9.37** | 3.54 | 1.56 | 1.37 | 2.91* | 2.85** | 0.38 | 2.75** |
| | | F | 328.95 ± 22.52 | 412.69 ± 37.02 | 402.03 ± 22.23 | | | | 1.71 | 1.54 | 0.22 | | | |

**Table S1**. Statistical analysis of the effects of MD lesions on cFos density across ACC and PrL cortical layers, using Two-way ANOVA comparison followed by the *post-hoc* analysis using Holm-Šídák (t) procedure. ACC: Anterior cingulate cortex; PrL: prelimbic cortex; Lc: lesion condition; MDmc: the medial and central subdivisions of the mediodorsal nucleus; MDl: the lateral subdivision of the mediodorsal nucleus. t1: MD sham vs. MDmc, t2: MD Sham vs. MDl; t3: MDmc vs. MDl; t4: male MD sham vs. female MD sham; t5: male MDmc vs. female MDmc; t6: male MDl vs. female MDl; M: male; F: female. Results are represented as mean (± SEM). *p < 0.05, **p < 0.01, ***p < 0.001.

| | | | CFos/PV Density (cFos+/PV+ cells/mm$^2$) (**Mean ± SEM**) LC groups | | | Two-way ANOVA | | | Pairwise comparisons within MD LC | | | Pairwise comparisons within sex | | |
|---|---|---|---|---|---|---|---|---|---|---|---|---|---|---|
| PFC sub-regions | Layers | Sex | MD sham | MDmc | MDl | LC $F_{(2,37)}$ | Sex $F_{(1,37)}$ | LC x Sex $F_{(2,37)}$ | t1 | t2 | t3 | t4 | t5 | t6 |
| ACC | 1 | M | 0 ± 0 | 0 ± 0 | 0 ± 0 | - | - | - | -- | -- | -- | -- | -- | -- |
| | | F | 0 ± 0 | 0 ± 0 | 0 ± 0 | | | | -- | -- | -- | | | |
| | 2/3 | M | 45.17 ± 3.71 | 31.79 ± 1.71 | 24.13 ± 3.69 | 31.89*** | 12.34** | 3.05 | 2.57* | 3.71** | 1.35 | 3.86*** | 1.87 | 0.36 |
| | | F | 65.86 ± 7.47 | 40.81 ± 2.96 | 26.13 ± 2.42 | | | | 4.54*** | 7.59*** | 2.90** | | | |
| | 5 | M | 45.71 ± 3.32 | 38.29 ± 3.22 | 45.71 ± 3.32 | 9.98*** | 1.19 | 9.34*** | 1.15 | 1.02 $e^{-015}$ | 1.10 | 3.42** | 1.15 | 2.63* |
| | | F | 67.88 ± 8.54 | 41.91 ± 5.32 | 27.96 ± 1.85 | | | | 3.42** | 6.28*** | 2.75** | | | |
| | 6 | M | 17.78 ± 2.71 | 17.83 ± 2.35 | 21.57 ± 4.58 | 0.14 | 7.32* | 0.56 | 0.01 | 0.80 | 0.82 | 2.24* | 1.72 | 0.73 |
| | | F | 26.68 ± 2.99 | 24.85 ± 3.99 | 24.89 ± 2.79 | | | | 0.56 | 0.68 | 0.10 | | | |
| PrL | 1 | M | 0 ± 0 | 0 ± 0 | 0 ± 0 | - | - | - | -- | -- | -- | -- | -- | -- |
| | | F | 0 ± 0 | 0 ± 0 | 0 ± 0 | | | | -- | -- | -- | | | |
| | 2/3 | M | 44.45 ± 3.87 | 25.48 ± 2.35 | 39.37 ± 6.65 | 11.36*** | 3.87 | 1.08 | 3,10* | 0.77 | 2.18 | 2.01 | 1.44 | 0.02 |
| | | F | 57.14 ± 4.92 | 34.28 ± 4.91 | 39.20 ± 3.92 | | | | 3.62** | 2.93* | 0.80 | | | |
| | 5 | M | 46.99 ± 3.87 | 26.96 ± 2.40 | 45.73 ± 8.11 | 27.80*** | 0.27 | 6.69** | 3.33** | 0.19 | 2.98* | 2.65* | 1.31 | 2.20* |
| | | F | 63.50 ± 5.67 | 19.05 ± 2.76 | 31.85 ± 2.66 | | | | 7.15*** | 5.5*** | 2.13* | | | |
| | 6 | M | 20.31 ± 3.18 | 21.37 ± 2.67 | 17.79 ± 3.77 | 0.15 | 0.40 | 0.70 | 0.23 | 0.52 | 0.76 | 0.27 | 0.04 | 1.31 |
| | | F | 19.04 ± 2.76 | 21.58 ± 3.87 | 23.91 ± 3.34 | | | | 0.54 | 1.08 | 0.52 | | | |

**Table S2**. Statistical analysis of the effects of MD lesions on CFos/PV density across ACC and PrL cortical layers, using Two-way ANOVA comparison followed by the *post-hoc* analysis using Holm-Šídák (t) procedure. ACC: Anterior cingulate cortex; PrL: prelimbic cortex; Lc: lesion condition; MDmc: the medial and central subdivisions of the mediodorsal nucleus; MDl: the lateral subdivision of the mediodorsal nucleus. t1: MD sham vs. MDmc, t2: MD Sham vs. MDl; t3: MDmc vs. MDl; t4: male MD sham vs. female MD sham; t5: male MDmc vs. female MDmc; t6: male MDl vs. female MDl; M: male; F: female. Results are represented as mean (± SEM). *p < 0.05, **p < 0.01, ***p < 0.001.

| | | | cFos/SOM Density (cFos+/SOM+ cells/mm$^2$) (**Mean ± SEM**) LC groups | | | Two-way ANOVA | | | Pairwise comparisons within MD LC | | | Pairwise comparisons within sex | | |
|---|---|---|---|---|---|---|---|---|---|---|---|---|---|---|
| PFC sub-regions | Layers | Sex | MD sham | MDmc | MDl | LC $F_{(2,37)}$ | Sex $F_{(1,37)}$ | LC x Sex $F_{(2,37)}$ | t1 | t2 | t3 | t4 | t5 | t6 |
| ACC | 1 | M | 0 ± 0 | 0 ± 0 | 0 ± 0 | - | - | - | -- | -- | -- | -- | -- | -- |
| | | F | 0 ± 0 | 0 ± 0 | 0 ± 0 | | | | -- | -- | -- | | | |
| | 2/3 | M | 29.20 ± 3.07 | 15.54 ± 2.51 | 29.20 ± 4.58 | 4.35* | 5.16* | 3.88* | 3.34***** | 8.31 $e^{-016}$ | 3.29****** | 1.85 | 0.87 | 2.87****** |
| | | F | 21.58 ± 1.98 | 19.04 ± 2.76 | 17.32 ± 2.38 | | | | 0.61 | 1.07 | 0.44 | | | |
| | 5 | M | 21.59 ± 3.07 | 26.66 ± 3.92 | 22.85 ± 4.40 | 1.49 | 2.03 | 3.84* | 1.15 | 0.26 | 0.83 | 1.39 | 2.31 | 1.59 |
| | | F | 27.94 ± 3.18 | 16.49 ± 1.98 | 15.57 ± 2.07 | | | | 2.51***** | 2.81****** | 0.21 | | | |
| | 6 | M | 19.05 ± 2.75 | 19.36 ± 3.77 | 22.87 ± 2.78 | 0.55 | 0.65 | 0.05 | 0.07 | 0.84 | 0.79 | 0.29 | 0.37 | 0.72 |
| | | F | 17.77 ± 2.71 | 17.79 ± 3.18 | 19.71 ± 2.48 | | | | 0 | 0.45 | 0.45 | | | |
| PrL | 1 | M | 0 ± 0 | 0 ± 0 | 0 ± 0 | - | - | - | -- | -- | -- | -- | -- | -- |
| | | F | 0 ± 0 | 0 ± 0 | 0 ± 0 | | | | -- | -- | -- | | | |
| | 2/3 | M | 24.12 ± 3.07 | 19.53 ± 2.43 | 16.52 ± 3.64 | 4.36* | 2.79 | 0.36 | 1.00 | 1.54 | 0.62 | 1.60 | 0.44 | 0.82 |
| | | F | 31.74 ± 3.87 | 21.59 ± 3.07 | 20.49 ± 3.72 | | | | 2.13 | 2.44 | 0.24 | | | |
| | 5 | M | 21.59 ± 1.97 | 18.58 ± 2.44 | 16.50 ± 3.64 | 1.48 | 0.50 | 0.07 | 0.69 | 1.09 | 0.45 | 0.57 | 0.11 | 0.52 |
| | | F | 24.13 ± 3.87 | 19.05 ± 2.76 | 18.87 ± 3.59 | | | | 1.14 | 1.21 | 0.03 | | | |
| | 6 | M | 19.05 ± 2.75 | 18.41 ± 3.17 | 19.05 ± 3.26 | 0.14 | 0.63 | 0.30 | 0.15 | 7.95 $e^{-016}$ | 0.14 | 0.88 | 0.15 | 0.62 |
| | | F | 15.23 ± 3.32 | 19.04 ± 2.76 | 16.35 ± 2.67 | | | | 0.88 | 0.26 | 0.64 | | | |

**Table S3**. Statistical analysis of the effects of MD lesions on cFos/SOM density across ACC and PrL cortical layers, using Two-way ANOVA comparison followed by the *post-hoc* analysis using Holm-Šídák (t) procedure. ACC: Anterior cingulate cortex; PrL: prelimbic cortex; Lc: lesion condition; MDmc: the medial and central subdivisions of the mediodorsal nucleus; MDl: the lateral subdivision of the mediodorsal nucleus. t1: MD sham vs. MDmc, t2: MD Sham vs. MDl; t3: MDmc vs. MDl; t4: male MD sham vs. female MD sham; t5: male MDmc vs. female MDmc; t6: male MDl vs. female MDl; M: male; F: female. Results are represented as mean (± SEM). *p < 0.05, **p < 0.01, ***p < 0.001.

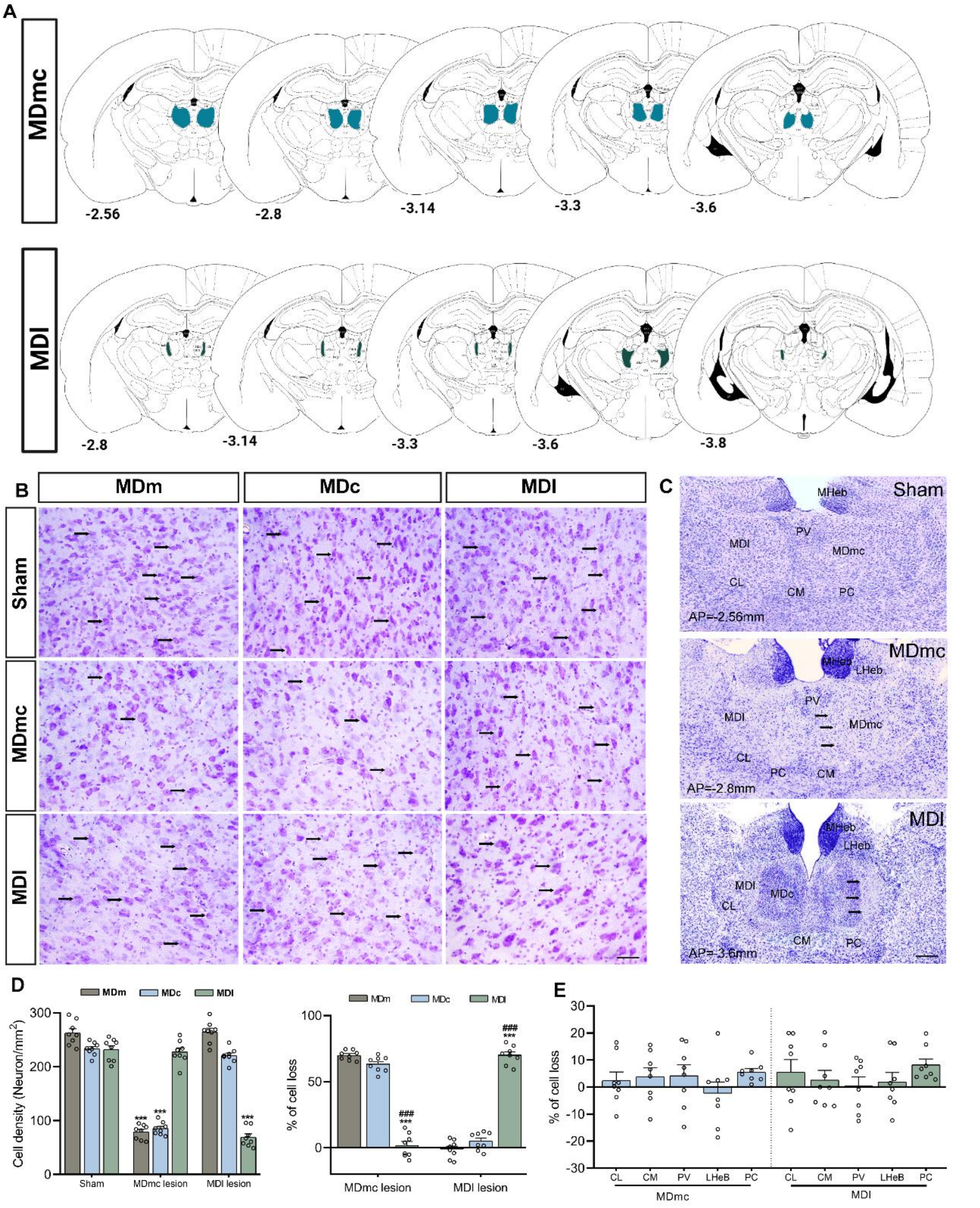


**Figure S1. Histological verification and quantification of subdivision-selective excitotoxic lesions in the mediodorsal thalamus (MDmc vs MDl).** (A) Schematic reconstructions of the maximal lesion spread (colored overlays) across rostro-caudal anteroposterior (AP) coordinates (mm, relative to bregma) for the MDmc (top row) and MDl (bottom row) lesion groups. Colored regions indicate the estimated area of neuronal loss within the targeted MD subdivision across animals. (B) High-magnification Nissl-stained micrographs images illustrating cellular architecture within MDm, MDc, and MDl for Sham, MDmc lesion, and MDl lesion animals. Black arrowheads indicate examples of Nissl-stained neuronal profiles used for density quantification (and highlight the reduction of identifiable neuronal somata in the targeted subdivision). Scale bar: 100µm. (C) Low-magnification thalamic sections showing the global lesion site and extent at selected AP levels for Sham, MDmc lesion, and MDl lesion groups. Scale bar: 300µm. (D) Quantification of neuronal density and lesion selectivity within MD subdivisions; Left: Neuronal cell density (neurons/mm²) in MDm, MDc, and MDl across groups; Right: Corresponding percent cell loss (normalized to Sham) for each lesion group, highlighting preferential loss in the targeted subdivision while sparing non-targeted MD regions. (E) Percent cell loss (relative to Sham) in surrounding thalamic nuclei to evaluate lesion specificity. Data are presented as mean ± SEM (n = 8). Data analyzed using one-way ANOVA followed by Holm-Šidák multiple-comparison *post-hoc* tests or t-tests. (***p < 0.001 Lesioned subdivision vs. Sham subdivision; ###p < 0.001 MDmc vs MDl). CL, centrolateral; CM, centromedial; PV, paraventricular; LHeb, lateral habenula; PC, paracentral.

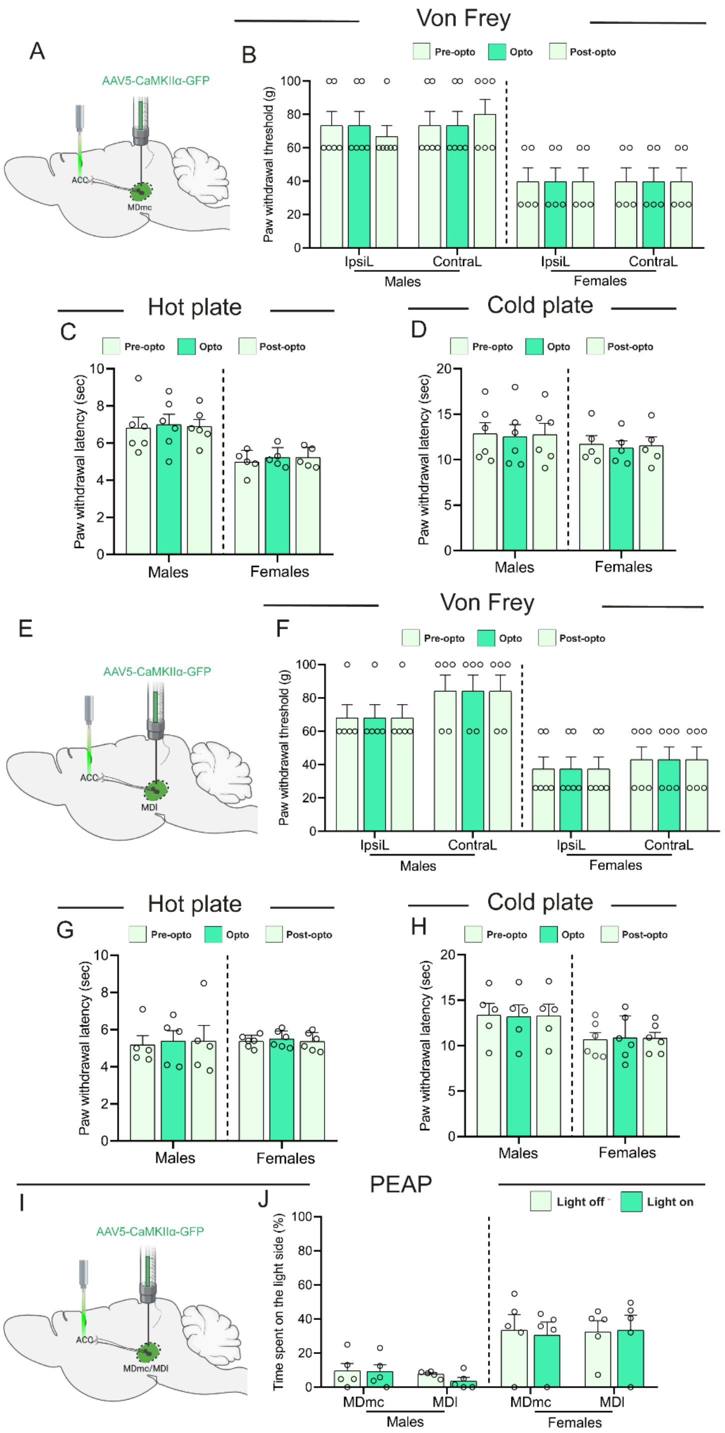


**Figure S2. GFP control experiments show that optical stimulation of MDmc/MDl-ACC pathways does not alter mechanical, thermal, or affective pain behaviors in male or female rats.** (A) Schematic illustration of the control experiment in which AAV5-CaMKIIα-GFP (without opsin) was injected into the MDmc and an optic fibre was implanted over the ACC. (B) paw-withdrawal thresholds (g) of the ipsilateral (IpsiL) and contralateral (ContraL) hind paws in the Von Frey test. (C,D) Hot-plate (C) and cold-plate (D) tests for the same MDmc-ACC GFP cohort, showing paw-withdrawal latencies (s). (E) Schematic of the analogous control experiment with AAV5-CaMKIIα-GFP injection into the MDl and an optic fibre over the ACC. (F) paw-withdrawal thresholds (g) in the Von Frey test. (G, H) Hot-plate (G) and cold-plate (H) paw-withdrawal latencies. (I) Schematic of the place escape/avoidance paradigm (PEAP) control experiment in which GFP-only virus was injected into MDmc or MDl, and an optic fibre was placed over the ACC. (J) Time spent on the light side (%) during PEAP. Data are presented as mean ± SEM. Sample sizes were as follows: MDmc-ACC-GFP (males n = 6, females n = 5) and MDl-PrL-GFP (males n = 5, females n = 6). For Von Frey, hot-plate, and cold-plate tests, repeated-measures two-way ANOVA was performed with stage and sex as factors. For PEAP, three-way ANOVA with factors subdivision and light condition and sex was used.

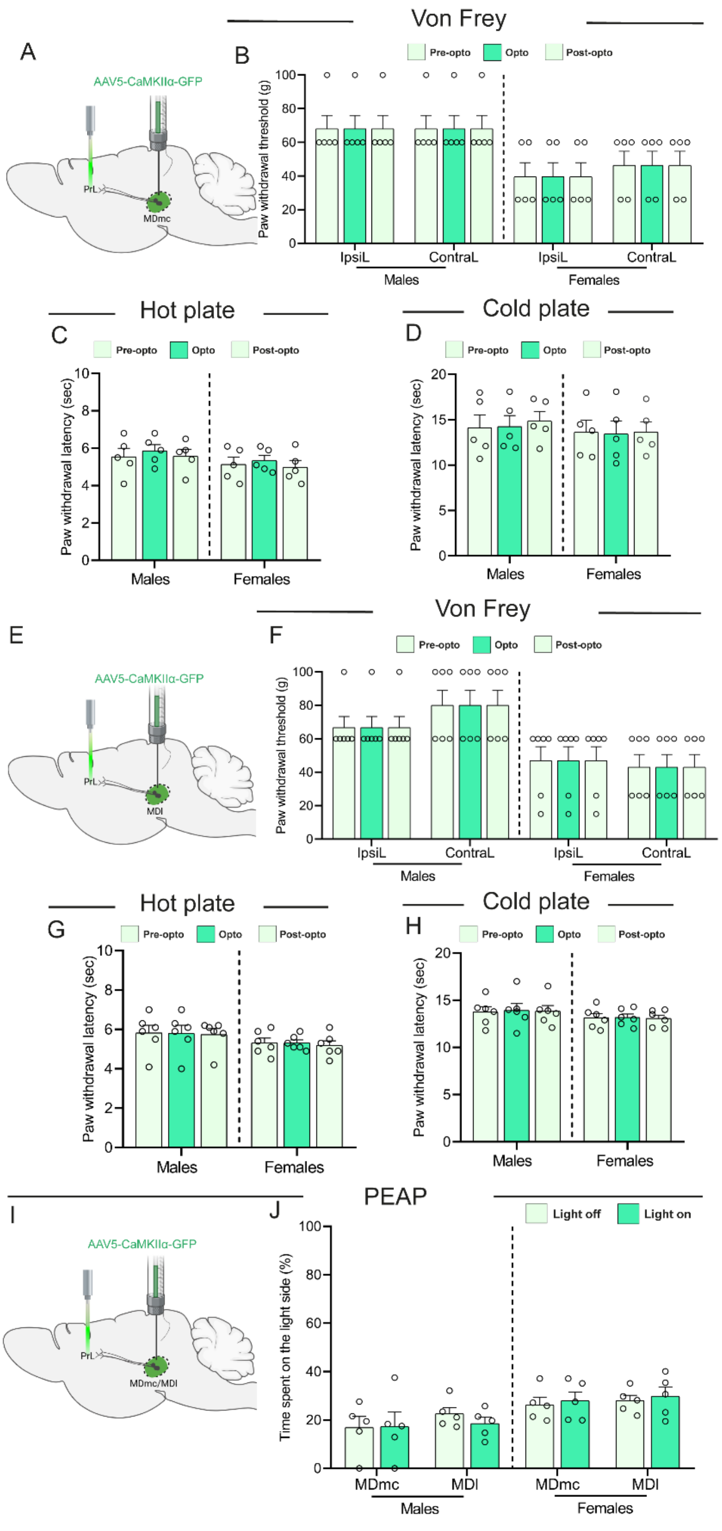


**Figure S3. GFP control experiments show that optical stimulation of MDmc/MDl-PrL pathways does not alter mechanical, thermal, or affective pain behaviors in male or female rats.** (A) Schematic illustration of the control experiment in which AAV5-CaMKIIα-GFP (without opsin) was injected into the MDmc and an optic fibre was implanted over the PrL. (B) paw-withdrawal thresholds (g) of the ipsilateral (IpsiL) and contralateral (ContraL) hind paws in the Von Frey test. (C,D) Hot-plate (C) and cold-plate (D) tests for the same MDmc-PrL GFP cohort, showing paw-withdrawal latencies (s). (E) Schematic of the analogous control experiment with AAV5-CaMKIIα-GFP injection into the MDl and an optic fibre over the PrL. (F) paw-withdrawal thresholds (g) in the Von Frey test. (G, H) Hot-plate (G) and cold-plate (H) paw-withdrawal latencies. (I) Schematic of PEAP control experiment in which GFP-only virus was injected into MDmc or MDl, and an optic fibre was placed over the PrL. (J) Time spent on the light side (%) during PEAP. Data are presented as mean ± SEM. Sample sizes were as follows: MDmc-PrL-GFP (males n = 5, females n = 5) and MDl-PrL-GFP (males n = 6, females n = 6).. For Von Frey, hot-plate, and cold-plate tests, repeated-measures two-way ANOVA was performed with stage and sex as factors. For PEAP, three-way ANOVA with factors subdivision and light condition and sex was used.

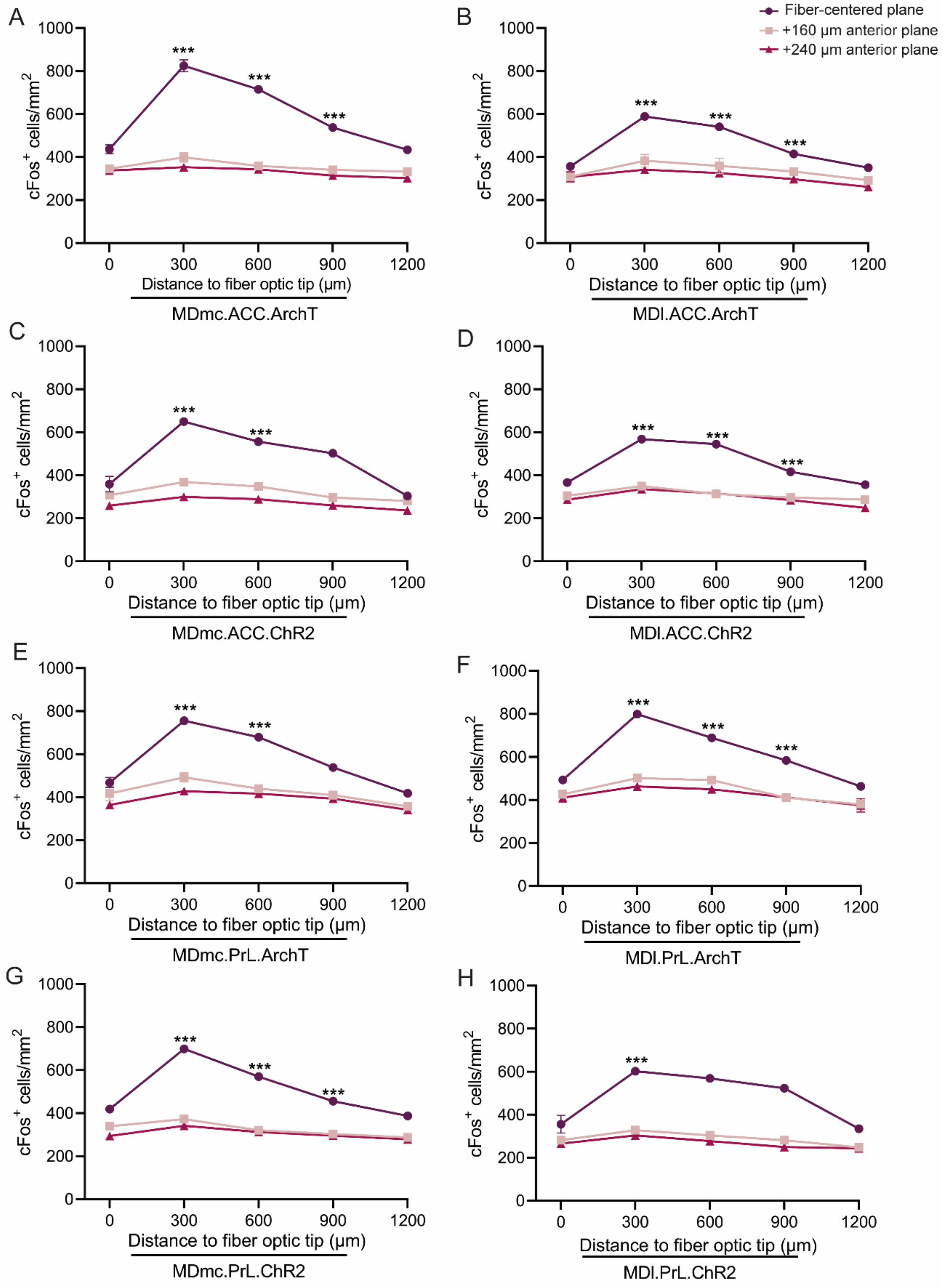


**Figure S4. Spatial mapping of the optogenetically driven cFos response around the fiber-optic tip in ACC and PrL cortex following MD subdivision-specific terminal modulation.** cFos positive cell density (cells/mm²) was quantified as a function of distance from the fiber-optic tip (0-1200 µm; pooled in 300-µm distance bins). Across conditions, cFos modulation was maximal in proximity to the fiber tip and progressively decayed with distance, indicating a spatially restricted optogenetic footprint. The ACC panels depict the distance-dependent cFos profile produced by MDmc-ACC and MDl-ACC terminal photoinhibition (ArchT) (A-B) and photoactivation (ChR2) (C-D) (separate plots for each MD subdivision and opsin). The PrL panels show the corresponding distance-dependent cFos profiles for MDmc-PrL and MDl-PrL terminal photoinhibition (ArchT) (E-F) and photoactivation (ChR2) (G-H). Data are presented as mean ± SEM (n = 4 per group) and were analyzed using a one-way ANOVA followed by Holm-Šidák *post-hoc* tests (*p < 0.05; **p < 0.01; ***p < 0.001 relative to the data point at 1200 µm).
